# SonoPrint: Acoustically Assisted Volumetric 3D Printing for Composites

**DOI:** 10.1101/2023.08.07.552292

**Authors:** Prajwal Agrawal, Shengyang Zhuang, Simon Dreher, Sarthak Mitter, Daniel Ahmed

## Abstract

Advancements in additive manufacturing in composites have transformed various fields in aerospace, medical devices, tissue engineering, and electronics, enabling fine-tuning material properties by reinforcing internal particles and adjusting their type, orientation, and volume fraction. This capability opens new possibilities for tailoring materials to specific applications and optimizing the performance of 3D-printed objects. Existing reinforcement strategies are restricted to pattern types, alignment areas, and particle characteristics. Alternatively, acoustics provide versatility by controlling particles independent of their size, geometry, and charge and can create intricate pattern formations. Despite the potential of acoustics in most 3D printing, limitation arises from the scattering of the acoustic field between the polymerized hard layers and the unpolymerized resin, leading to undesirable patterning formation. However, this challenge can be addressed by adopting a novel approach that involves simultaneous reinforcement and printing the entire structure. Here, we present SonoPrint, an acoustically-assisted volumetric 3D printer that produces mechanically tunable composite geometries by patterning reinforcement microparticles within the fabricated structure. SonoPrint creates a standing wave field that produces a targeted particle motif in the photosensitive resin while simultaneously printing the object in just a few minutes. We have also demonstrated various patterning configurations such as lines, radial lines, circles, rhombuses, quadrilaterals, and hexagons using microscopic particles such as glass, metal, and polystyrene particles. Furthermore, we fabricated diverse composites using different resins, achieving 87 microns feature size. We have shown that the printed structure with patterned microparticles increased their tensile and compression strength by ∼38% and ∼75%, respectively.

## Main

Recent years have seen a growing interest in additive manufacturing of composite structures due to their high strength-to-weight ratio, corrosion resistance, and design flexibility, features which make composites attractive for industries such as aerospace (1–3), automobiles (4), construction (5–7), medicine (8, 9), and more. “Composite” refers to a structure constructed from two or more materials: a continuous phase, the matrix, and a discontinuous phase, the reinforcement, that lends strength to the structure (10). Alignment of the reinforcement in a composite can significantly improve the structure’s mechanical properties, i.e., its strength, stiffness, and toughness, by creating more efficient load transfer pathways and reducing stress concentrations between the matrix and reinforcement (11). Thus, composite structures offer improved performance. For example, composites reinforced with carbon nanotubes (CNTs) exhibit a 200% increase in Young’s modulus (12), while aligning microfibers in a 3D printed object resulted in 77% more strength compared to randomly oriented fibers (13). Current methods for aligned reinforcement involve passive or active forces (14–18). Active approaches rely on external energy sources such as electric, magnetic, or acoustic fields to pattern desired microparticle motifs; these methods offer higher control and precision than passive approaches that use shear (19, 20), gravitational (21), hydrodynamic (22, 23), and other such forces for alignment (24). Active methods are able to correct manufacturing variations, such as uneven thicknesses or warpage (25, 26); this promotes more precise alignment and greater flexibility, and hence enhances the efficiency of the manufacturing process.

Although electric (14, 27–33) and magnetic (15, 34–38) field-assisted reinforcement strategies have been applied to 3D printing of composites, both rely on intrinsic properties of the particles, such as surface charge and magnetism, and hence these strategies are limited in choice of materials. In addition, they are limited in generating complex reinforcement patterns. Moreover, electric and magnetic alignment methods require complex experimental setups, and are relatively expensive (14, 16, 34). Acoustics offer an attractive alternative modality for manipulating microscopic particles within a host resin, as the effect of sound on particles is independent of features such as size, geometry, and charge (39–41). However, incorporating acoustics into most conventional 3D printing methods presents a significant challenge. As each layer is printed, a complex scattered acoustic field arises between the polymerized hard layers and the unpolymerized resin just above/below it. Thus, control over the internal patterning of microparticles become inadequate in most extant acoustically-assisted 3D printing methods (18, 42–44).

The recently developed vat-polymerization-based manufacturing technology, termed volumetric printing, solves this problem by printing structures as whole (45). Derived from the mathematical models of computed tomography (CT) and intensity modulated radiation therapy (IMRT) scans, this approach can print 3D structures as whole volumes, resulting in a faster build time compared to layer-by-layer additive manufacturing (45, 46). However, current volumetric printing methods are limited to high-viscosity resins (4-11 Pa.s), while existing acoustically-assisted printing strategies implemented in extrusion (17, 25, 47–50) and vat-polymerization (18, 42–44, 51) methods employ low-viscosity resins (0.1-5 Pa.s). The more limited transmission of acoustic waves through high-viscosity resins (18, 42, 52) presents its own difficulty for acoustic patterning of particles. To date, no acoustically-assisted volumetric printing techniques have been demonstrated.

Here, we introduce SonoPrint, an acoustically-assisted volumetric 3D printing technique that uses acoustic energy to trap microparticles in a volumetrically-fabricated 3D composite structure. In developing this technique, we first constructed volumetric- and acoustic-compatible experimental setups consisting of piezoelectric transducers (PZTs), index-matching liquid, and a vat filled with resin (matrix) and microparticles (reinforcement). The reinforcement consisted of various materials such as glass, polystyrene, hollow materials, and metals in sizes from 1 to 250 microns in di-pentaerythritol pentaacrylate (PETA) and aliphatic urethane diacrylate (UDMA) resins. Reinforcements were aligned in the resin using various numbers of PZTs with different placement configurations and different frequencies, allowing patterning in the form of parallel lines, radial lines, circles, rhombuses, quadrilaterals, hexagons, and polygons. The size, thickness, and distance between pattern units were controllable based on the PZT setup; for example, we produced lines separated by 0.46 to 0.96 mm. Subsequently, through volumetric printing, we were able to incorporate these patterned microparticles into a composite geometry within minutes. Optimizing the parameters yielded composite structures with a feature size of around 87 µm. When observed under a microscope, the printed structures displayed consistent reinforcement throughout the structure. When tested for mechanical strength using tensile and compression tests, the strength of structures with aligned microparticles increased by ∼38 and ∼75% respectively. Accordingly, we expect our proposed composite fabrication method to open up a whole new realm of applications for volumetric printing.

### Experimental Setup

SonoPrint comprises several components, including a projector, a glass resin vat, a vat rotation mechanism, an acoustic setup, a function generator, an amplifier, a control unit, and the SonoPrint software. The projector displays images generated by the software onto the rotating resin vat, which contains the photosensitive liquid resin and microparticles, as shown in **Fig. 1*A***. The vat is enveloped by an index-matching liquid. The acoustic setup consists of PZTs positioned around the resin container; these are controlled by a function generator and an amplifier to produce an acoustic wave field. The resonance frequencies of the PZTs were measured using an oscilloscope. **Fig. 1*A*** and ***B*** depict the SonoPrint system components alongside a model geometry they were used to fabricate.

**Figure 1.**
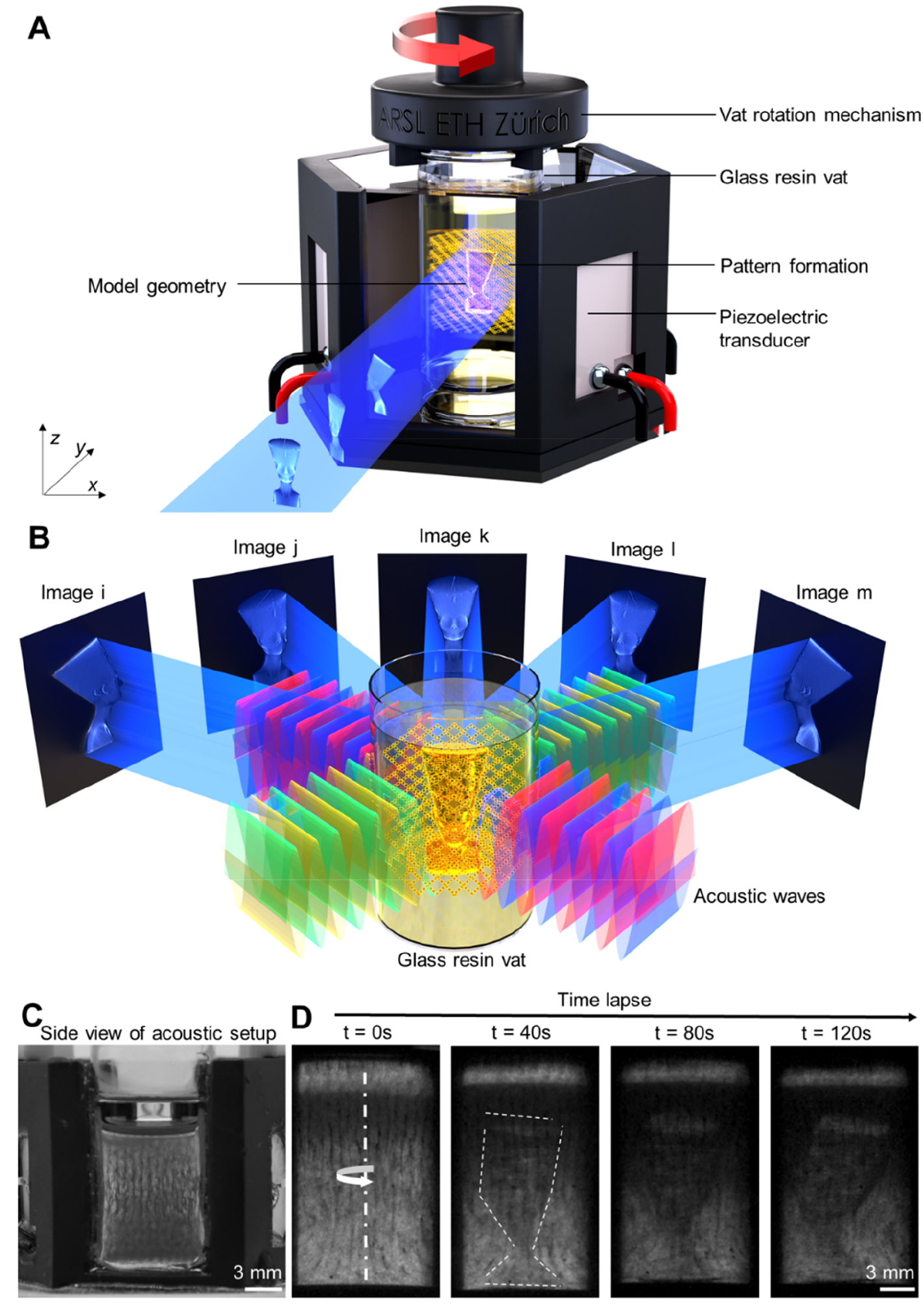
SonoPrint: An acoustically-assisted volumetric 3D printer. **A.** Schematic illustrating the setup of a SonoPrint printer. Images of Nefertiti model geometry are being projected on a rotating glass resin vat (hereafter termed as vat) focused at the center of the vat. The vat contains photopolymerizable resin with acoustically-aligned microparticles. A four piezoelectric transducer (PZT)*−*setup is used for the patterning of microparticles. A Nefertiti model geometry is also formed within the vat surrounded by index-matching liquid. **B.** The schematic illustrates the generation of bulk acoustic waves by the PZTs and their transmission through the vat to pattern the microparticles inside the resin. Image projections from i to m illustrate the projections of various images generated by volumetric software that are projected at different angles and at their corresponding times. As shown in the schematics, this results in the fabrication of a composite model geometry of Nefertiti with patterned microparticles in a desired manner. **C.** Images show side view of patterned microparticles in the four*−*PZT acoustic setup when all four PZTs are excited at 1.01 MHz, 30 V_PP_ and 25 °C. Scale bar 3mm. **D.** This figure illustrates the fabrication of 3D Nefertiti composite geometry over a time span of 120 seconds with microparticle pattern obtained in C, also see **Movie S6**. Scale bar: 3 mm.

When patterning microparticles within the vat, we employed three acoustic setups, each equipped with two, four, and six PZTs, respectively. The two*−* and four*−*PZT acoustic setups were made compatible with volumetric printing by incorporating glass panels on the front and back, enabling the transmission of projected light. The other sides of these two setups housed the PZTs, which generated bulk acoustic waves (see **Fig. 1*B*** showing the four*−*PZT setup). Within each setup was placed a transparent glass vat, with a refractive index (RI) of 1.516; this was surrounded by glycerol with RI of 1.4731 (53). These index-matching liquids had negligible impact on acoustic transmission, with acoustic attenuation loss of 0.5% for glycerol; see also **Supporting Text 1**. Glycerol was used as the index-matching liquid for experiments due to it better refractive index matching with the resins and the setup. The PZTs were connected to a function generator and amplifier to generate a standing acoustic wavefield in the vat to align the microparticles, see **Fig. 1*C***. In the six*−*PZT setup, no light can pass through because all six sides are housed with PZTs (see **Fig. 2*A*** and ***B***), so after achieving the desired microparticle pattern, the vat was transferred to a square container made of glass from all four sides to allow volumetric printing, see **Fig. S7*A*** and ***B***.

**Figure 2.**
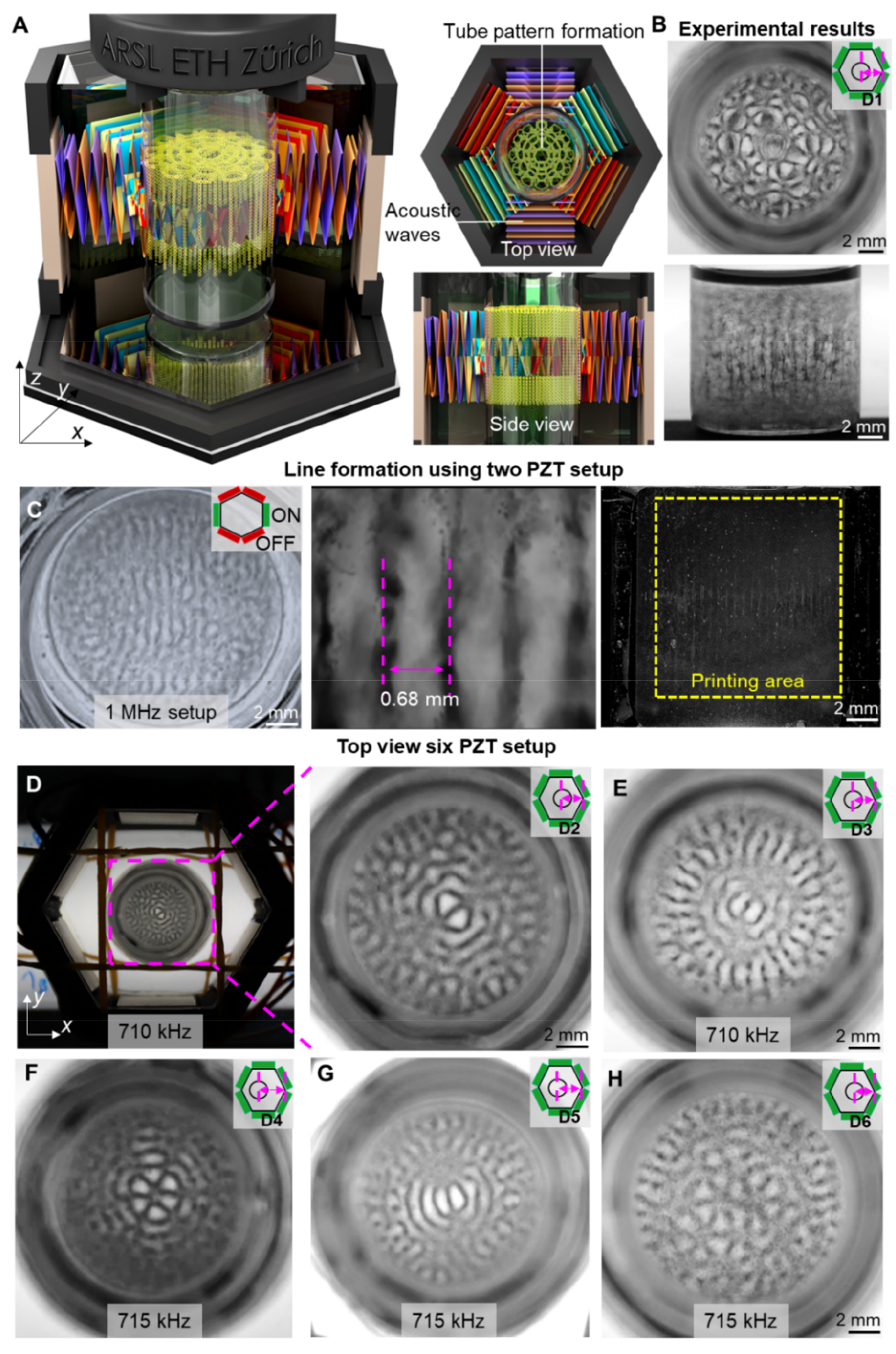
Acoustic patterning of microparticles. **A.** A schematic representation of a six piezoelectric transducer (PZT)*−*acoustic setup that utilizes six identical PZTs orthogonally positioned in *xz*-plane being actuated simultaneously at resonance frequency to produce bulk acoustic waves for patterning microparticles in the desired manner. A top and side view of the setup also illustrates the patterns from these views. **B.** Displays the top and side view of the six*−*PZT setup when the glass resin vat (hereafter termed vat) is placed at the center of the acoustic setup and all six PZT are actuated at 715 kHz, 30 V_PP_ and 25 °C. It shows the patterns as generated in illustration in A. D1, corresponds to ∼21 mm distance from the extreme corner of the vat. Scale bar: 2 mm. **C.** Results uses 1 MHz two*−*PZT acoustic setup, excited at their resonance frequency of 1.02 MHz, at 30 V_PP_, and 25 °C. The images show their top, bottom-microscopic, and side views. 5-50 micron soda lime glass microspheres were used in di-pentaerythritol pentaacrylate resin (PETA). Scale bar: 2 mm for side and top view. **D.** Illustrates the six*−*PZT acoustic setup. The zoomed in view shows the vat located at D2, i.e., ∼20.1 mm distance in the *x*-axis, excited at a frequency of 710 kHz, an amplitude of 30 V_PP_, and a resin temperature of 25 °C. Scale bar: 4 mm. **E.** Image shows microparticle pattern generated in the six*−*PZT setup by moving the vat along the *x*-axis by D3, i.e., ∼19.3 mm from the extreme corner of the acoustic setup and using a frequency of 710 kHz, an amplitude of 30 V_PP_, and a temperature of 25 °C. Scale bar: 2 mm. **F - H** A top view of the patterning is shown in relation to the change in placement of the vat container relative to the *x*-axis following excitation of all six PZTs in the six*−*PZT acoustic setup. Patterns were formed with stainless steel microspheres of 1-22 microns in PETA resin. A frequency of 720 kHz, amplitude of 30 V_PP_, and resin temperature of 25 °C was employed. The glass resin vat was moved along *x*-axis at distance D4 – D6, corresponding to ∼25.2, ∼23.1, and ∼19.3 mm distances from the extreme corner of the setup. Scale bar: 2mm.

In our study of microparticle patterning, we employed polystyrene microparticles (15 µm), hollow glass microspheres (*∼*49 µm), soda-lime glass microspheres (*∼*22, *∼*113, and *∼*275 µm), and stainless steel metal microspheres (*∼*11 and *∼*101 µm). When exposed to an acoustic wavefield, the microparticles were subjected to acoustic radiation forces. We estimated the primary radiation force acting on solid microparticles using the gradient of the Gor’kov potential; see also **Supporting Text 1**. To determine the directionality of that force, we calculated the acoustic contrast factor Φ; we found this factor to be positive for all tested particle types, indicating movement towards the pressure nodal position (52, 54), an effect illustrated in **Fig. 1*B***. We further investigated the acoustic parameters, such as frequency and amplitude, to achieve desired microparticle patterns, see **Movie S1** and **S2** for line pattern using 1 MHz PZTs. Finally, we evaluated the compatibility of the SonoPrint printer system with resins of different viscosities: PETA (4-10 Pas) and UDMA (7-11 Pas).

Following acoustic patterning as seen in **Fig. 1*C***, we utilized volumetric printing to create the structures. In this process, the desired 3D model in standard triangle language (STL) is fed to the volumetric printing software, which generates a series of 2D images (45, 55, 56). Typically, one image is generated for each rotation angle, resulting in 360 images for a complete 360 degree rotation (45). The resulting images are projected onto the vat’s center, which contains the polymerizable resin and patterned microparticles. The vat is rotated during this process, ensuring each image is projected at the correct angle; as illustrated in **Fig. 1*B***. Polymerization is initiated upon reaching a significant energy threshold by the projected light, which usually takes 30 to 240 seconds resulting in the formation of 3D structures, as shown in **Fig. 1*D*** and **Movie S6**. After fabrication, the structures were removed from the resin vat, and cleaned (see Methods). We examined the pattern formations in the fabricated geometries. Finally, we assessed the feature size of the printed geometries using scanning electron microscopy (SEM) and mechanical strength of the structures using tensile and compression tests.

### Acoustic patterning of microparticles and pattern characterization

We began by using a two*−*PZT acoustic setup to create specific patterns, then advanced to more intricate ones requiring a four*−* and six*−*PZT setup. In the two*−*PZT configuration, a pair of parallel PZTs were placed opposite each other and with the simultaneous acoustic excitation at the same frequency and voltage, generated a standing acoustic wavefield within the resin vat as shown in **Fig. S4 *A*, *B*,** and ***C***. This wavefield consisted of a series of pressure nodes and antinodes, in which acoustic forces pushed the microparticles toward the pressure nodes, resulting in microparticle patterns in the *x*-direction and distributed as sheet in the *yz*-plane as seen in **Fig. S4*A***. In order to vary the distance between the formed line patterns, we used PZTs of different resonant frequencies. For example, for varying the distance between the formed line patterns from 0.45 to 0.96 mm we used PZTs with resonance frequencies of 710 kHz, 1 MHz, and 1.5 MHz. First, by using 1 MHz PZTs, we were able to achieve a distance of 0.68 mm between two formed line patterns, as seen in **Fig. 2*C***. Similarly, **Fig. S4*D*** and ***E*** depict line pattern formations with distance between two formed lines of 0.96 and 0.45 mm using two*−*PZT setup of 710 kHz and 1.5 MHz, respectively, as observed from the side, top, and bottom perspectives. Additionally, by adjusting the microparticle concentration, we achieved variation in the line thickness from ∼0.25 to ∼1 mm. **Movie S5** shows the varied line thicknesses achieved when using 20 to 80 milligrams of 5-50 micron soda lime glass microspheres in 20 grams of PETA. All patterns obtained in two*−*PZT setups were reproducible in four*−* and six*−*PZT setups by exciting any pair of parallel PZTs.

When creating line patterns, the resin temperature was maintained at 25 °C. As the temperature increases, it becomes easier to create patterns using sound waves; however, the resin becomes less viscous, and the microparticles tend to settle over time due to gravity, which adversely affects the printing. For example, when the resin temperature is at 45 °C, the patterned stainless steel microspheres (1-22 μm) do not remain intact and settles to the bottom in a few minutes. In order to keep the pattern intact during printing, the resin temperature was kept low. Nevertheless, when PZTs are operated at a high voltage for an extended period of time, they generate heat, which raises the temperature of the resin. When PZTs are operated at 60 V_PP_ for up to two minutes, the resin temperature can rise from 25 °C to as high as 50 °C. As a means of controlling the resin’s viscosity, we tested different temperatures to determine the duration for which the pattern, created through the application of sound waves, remained intact. We observed that when the resin temperature was maintained below 30°C, the pattern exhibited stability, persisting for over ten minutes, as illustrated in **Fig. S3**. We were able to achieve this desired temperature, by applying low voltages and pulsed excitations. As an example, to achieve the line pattern shown in **Fig. 2*C***, we applied 30 V_PP_ for 30 seconds while maintaining the resin temperature below 30 °C.

The four*−*PZT setup incorporates four orthogonally-positioned PZTs; activating these PZTs at a similar frequency produces rhombus patterns in the *xy*-plane, which were projected overall in the *z-*axis (**Figs. S6*A*, *B*,** and ***C***). To demonstrate the formation of rhombus patterns with different particles, we used 15 µm polystyrene particles in PETA resin as seen in **Fig. S6*E***. Using this setup, we also obtained circular, quadrilateral and other polygon patterning by exciting all four PZTs simultaneously at different frequencies, as seen in **Fig. S6*F***, and **Movie S3**. In addition, we investigated the effect of stimulating only two parallel PZTs, which resulted in orientated line patterns, see **Movie S4**. Patterns obtained from four*−*PZT setup were reproducible when two pairs of parallel PZTs were excited in the six*−*PZT setup.

As shown in **Figs. 2*A*** and ***B***, a tube-like pattern formation is generated by simultaneously activating all six PZTs at the same frequency in the six*−*PZT acoustic setup. The pattern in **Fig. 2*B*** was obtained by placing the vat in the center, using 715 kHz frequency, 30 V_PP_ amplitude, and 25 °C temperature. Additionally, we generated various patterns by placing the vat centered along the *y*-axis and moving it along the *x*-axis; **Fig. 2*D*** illustrates the six*−*PZT setup from top view. To create the polygonal pattern, we started with the vat located at D2, i.e., *∼*20.1 mm distance in the *x*-axis, a frequency of 710 kHz, an amplitude of 30 V_PP_, and a resin temperature of 25 °C. **Fig. 2*D*** illustrates the patterns formed with a zoomed-in view. A second pattern, as shown in **Fig. 2*E***, was obtained by using the same parameters and by just moving the vat along the *x*-axis by *∼*19.3 mm from the corner denoted by D3. Finally, we positioned the resin vat at three different positions, D4-D6 corresponding to *∼*25.2, *∼*23.1, and *∼*19.3 mm distances from the corner of the acoustic setup, respectively. With acoustic actuation at 720 kHz, 30 V_PP_, and 25 °C we obtained tubular patterns as shown in **Figs. 2*F*-*H***. **Supporting Text 2** and **Figs. S7** and **9** show other patterns achieved with the six*−*PZT setup. These findings demonstrate that with our setups, multiple pattern types can be developed, and versatile, dynamic modulation of patterns is feasible.

### Volumetric printing

Once the desired patterning has been achieved, the polymerization process usually takes between 30 and 240 seconds for printing structures measuring up to 4 cm in height. Throughout the fabrication process, the patterns achieved in the resin remained intact, which can be attributed to the high viscosity (4 to 11 Pas) of these resins, see **Fig. S3**.

For printing, initially, all PZTs in six*−*PZT acoustic configuration were excited at the same frequency in order to create a microparticle pattern. Using the volumetric fabrication process, the patterning was incorporated into the Nefertiti model geometry. The side and top views of the fabricated geometry at different planes and depths are shown in **Fig. 3*A*** and ***B***. Later, to assess the microparticle patterning consistency inside a 3D printed structure we printed a cylindrical geometry and observed the pattern at different depths and angles as seen in **Fig. S11*A*-*D***. In order to replicate the straight and radial line microparticle pattern, we printed two Nefertiti structures with 1-22 micron stainless steel microspheres mixed in PETA resin. We achieved the desired parallel and radial line patterns consistently in the whole Nefertiti geometry by patterning them as defined in **Supporting Text 2**, and then fabricating the structures using volumetric printing mechanism. The fabricated geometry and microparticle pattern are shown in **Fig. 3*C*** and ***D*** respectively. Additionally, a cylindrical structure was printed to demonstrate complex quadrilateral patterns to be achieved in a fabricated structure. The patterning was achieved by exciting four PZTs at 1.02 MHz, 30 V_PP_, and 25 °C. The top, side and zoomed-in top view in **Fig. 3*E***, shows the formed cylindrical structure with quadrilateral patterns.

**Figure 3.**
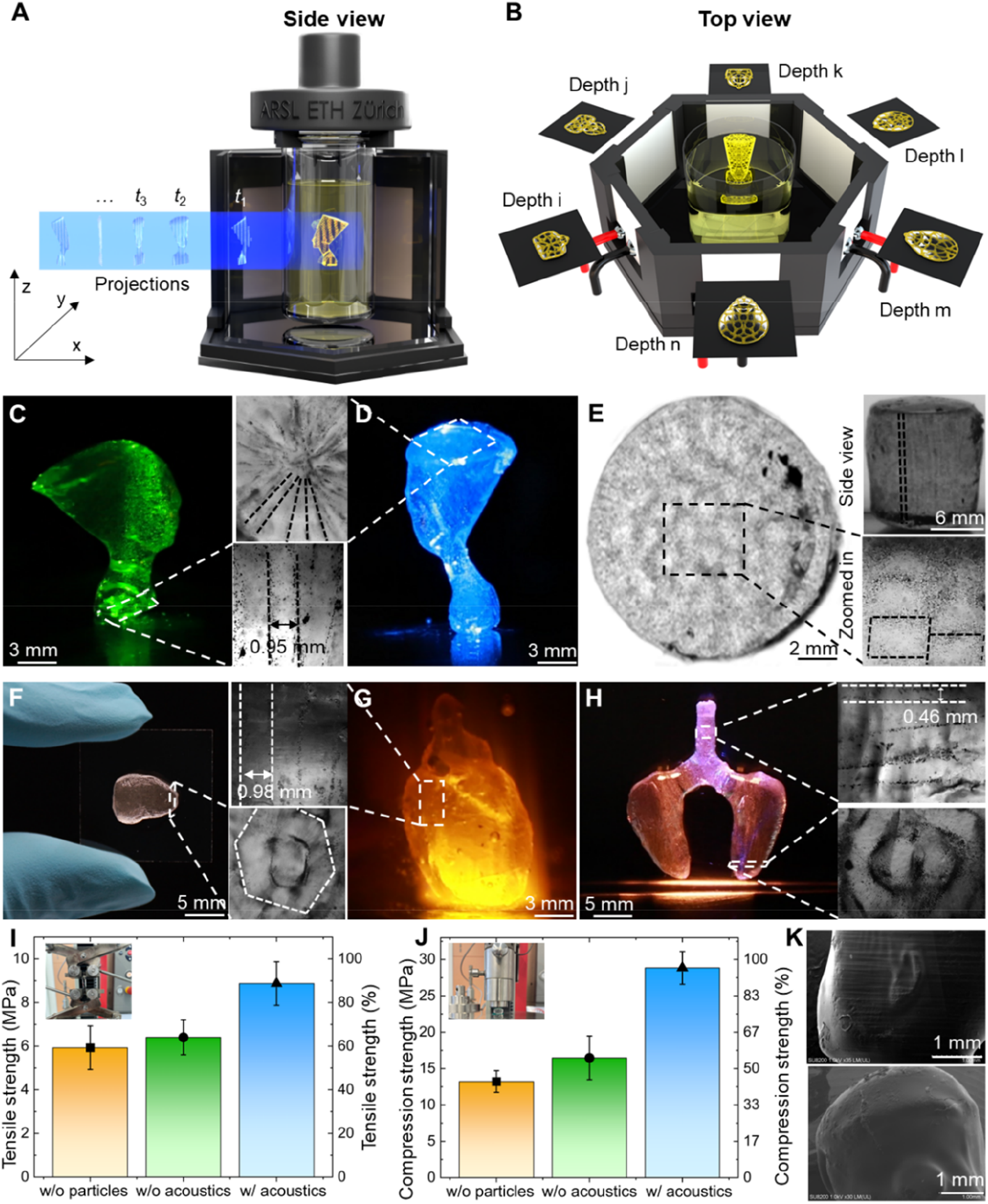
Analyses and characterization of the printed composite structures generated by the SonoPrint printer. **A** and **B.** A schematic of the fabricated Nefertiti model with patterning as shown from both a side and top perspective. **C** and **D.** The image displays a complex Nefertiti composite geometry printed with parallel and radial line patterns respectively. Patterns were generated with two*−*PZT setup at 1.02 MHz, 30 V_PP_, and 25 °C. Also, a microscopic zoomed-in view shows the patterning. Scale bar: 3 mm. **E.** The images illustrate a cylindrical composite geometry with patterns generated by exciting four PZTs simultaneously at 1.02 MHz, 30 V_PP_, and 30 °C. The top, bottom and zoomed-in top view are shown. Scale bar: 2 mm for left image and 6 mm top right corner image. **F.** Displays a miniature printed liver geometry when held on a glass slide. An enlarged zoomed view illustrates the hexagonal patterning inside the geometry that was obtained by the excitation of all six PZTs at 710 kHz, 30 V_PP_, and 25 °C. Scale bar: 5 mm. **G.** The image shows a miniaturized hollow heart geometry with line patterning obtained by excitation of a pair of parallel PZTs at a frequency of 715 kHz at 30 V_PP_ and 25 °C. A zoomed in view shows the line pattern with 5-50 micron soda line glass microspheres. Scale bar: 3 mm. **H.** The figure depicts a miniaturized lung geometry, printed in three parts, where the right and left lungs have polygonal microparticle pattern, while the trachea incorporates horizontal lines. Six and two PZTs were excited at 710 kHz, 30 V_PP_, and 25 °C in order to achieve polygon and line pattern respectively. A zoomed-in view illustrates the pattern. Scale bar: 5 mm. **I** and **J.** Displays a bar graph comparing the results of the tensile and compression tests, respectively. The test samples were printed without microparticles, with microparticles but without acoustic alignment, and with microparticles aligned vertically in tensile samples and horizontally in compression samples. An image of the testing setup can be seen in the top left corners. Error bar: 3 samples. **K.** Shows the SEM image of the Nefertiti and Mario composite geometry. Scale bar: 1mm.

To demonstrate complex printing with a variety of patterns, we reproduced miniaturized models of liver, heart, and lungs, with hexagonal, line, and polygon patterns. By stimulating all six PZTs simultaneously at 710 kHz, 30 V_PP_, and 25 °C, the microparticles were patterned in hexagonal shapes. Following this, the vat with aligned microparticles was placed in the volumetric printing setup, shown in **Fig. S7*A*** and ***B***. As a result, the liver model geometry was printed as shown in **Fig. 3*F***, and the resulted hexagonal pattern is shown in the zoomed-in view. To replicate a miniaturized hollow heart model, we formed line patterns with six*−*PZT setup using the stated above method for patterning. Here, 5-50-micron soda lime glass microspheres were patterned in rhodamine-mixed PETA resin. An image of the fabricated heart geometry is shown in **Fig. 3*G*** when a beam of blue light is shined using a microscope. The zoomed-in figure illustrates the line pattern. Lastly, a model of the lungs was printed in three parts and then assembled together. In the first step, 1-22 micron stainless steel microspheres were used in the printing of the left and right lungs in order to incorporate a polygonal microparticle pattern as obtained in **Fig. 2*D***. Alternatively, trachea was printed with horizontal line pattern. Following this, the geometry was assembled together as shown in **Fig. 3*H***. The zoomed-in image shows microparticle patterning in the lungs and trachea.

### Characterization, analysis, and applications of printed composite geometry

We conducted mechanical tests to evaluate the mechanical strength of SonoPrint fabricated structures with aligned microparticles compared with structures printed without microparticles or with microparticles but not aligned. The samples with microparticles contained 1-22 micron stainless steel metal microspheres. A 1.5 MHz two*−*PZT acoustic setup excited at 1.51 MHz, 30 V_PP_, and 25 °C was used for the alignment of particles. For tensile samples, the microparticles were aligned vertically, i.e., in parallel to the applied tensile force direction, whereas for compression samples, they were aligned horizontally, i.e., in perpendicular direction to the applied compression force. The tensile strength of the acoustically aligned samples was *∼*38% higher than the samples with microparticles that were not aligned (see **Fig. 3*I***). The compression test demonstrated that acoustically aligned samples demonstrated a *∼*75% increase when compared with structures with microparticles and without acoustical alignment (see **Fig. 3*J***). Thus, aligned microparticles appear to provide an advantage in terms of mechanical strength. Also, SEM microscopy was also used to verify that the structural features of the printed 3D geometry were not compromised. The feature size of Nefertiti’s and Mario’s composite structures was examined, as shown in **Fig. 3*K***.

## Discussion

SonoPrint comprises the first acoustically assisted volumetric 3D printing system. It obviates the limitations of fabricating composite structures with aligned reinforcement of particles. Using SonoPrint, we successfully demonstrated microparticle alignment into different shapes that were then integrated into various 3D structures. We printed complex geometries like Nefertiti, Mario, and anatomical organs such as the heart, lungs, and liver using aligned microparticles in radial lines, rhombus, parallel lines, hexagonal, and polygonal shapes. A consistent reinforcement patterning was achieved and shown throughout the structure. The fabricated composite structures possessed a feature size of around 87 µm. Also, tensile and compression strength of the structures with aligned microparticles was enhanced by ∼38 and ∼75% respectively.

In SonoPrint, a factor that affects print quality is the concentration of microparticles in the resin. A high number of microparticles makes it difficult for light to pass through the resin, thereby adversely affecting the print quality. We studied this effect by using varying amounts of 1-22 micron stainless steel microspheres in PETA resin, ranging from 5 to 50 milligrams per four grams of resin, as illustrated in **Fig. S10**. In the future by employing microparticles only in a specific region of the vat and not in the overall resin may help increase microparticle concentration in fabricated structures.

The SonoPrint additive manufacturing platform enhances control over properties in tailor-made 3D composites. The developed technology can foster developing new robotic materials for biohybrid robots and tissue constructs. The 3D printing of biocomposites provides opportunities in designing and developing innovative biohybrid robots, grippers, and pumps. Tissue constructs that can emulate specific biological tissues’ physiology and functionality through intricate cellular arrangements, holds significance in tissue engineering, drug screening, and beyond. Precisely organizing cardiomyocytes into parallel lines ensures cardiac constructs’ efficacy, while aligning endothelial cells is pivotal for functional vascular structures. Hexagonal patterns hold promise in building liver tissue constructs. Currently, the accurate replication of tissue cellular patterns is difficult; and this can be overcome using SonoPrint. We envision SonoPrint opening up new applications for volumetric printing in soft electronics, tissue fabrication, aviation, sportswear, and other fields.

## Materials and Methods

### Volumetric Printer Hardware

The hardware for volumetric printer consists of a rotating vat made of glass (ETH Internal D-Phys Shop part number 40053.1) onto which 2D image slices are projected, a light source (Si cube’s SM9 with 405 nm wavelength), to generate the projection images, a lens to focus (Edmus Optics, part number #67-187), a motor to control z-axis movement (Nema 17, Conrad order number: 1597325-AW), and an acoustic chamber which was fabricated using index-matching glass, self-cut into desired shape using glass slides (from Sigma Aldrich, part number: CLS294775X25) containing a index-matching liquid i.e., glycerol (from Sigma Aldrich part number G5516-100ML)) used for lesser light diffraction. The 2D slices generated by the software are projected at an intensity of ∼0.1 to 2.0 milli watt (mW)/cm^2^ at different angles but with consistent angular velocity onto the vat containing the photopolymerizable ink PETA or UDMA, which is rotated via a stepper motor (Conrad part number: 1597325-AW). Volumetric curing of resin is depicted in **Fig. 1*D*** and the fabricated 3D geometries are shown in **Fig. 3**. The vat container is confined in the acoustic chamber that generates the acoustic patterning initially. Local oxygen depletion is performed: an initial phase of free radical activation by light followed by rapid quenching, and then deactivation by oxygen. The acoustically patterned microparticles are also trapped inside the 3D geometry, making it a composite material with the patterned microparticles acting as reinforcement as shown in **Fig. 3**.

### Volumetric Printer Software

The software for the printer is based on the open-source code of Kelly et al.(45), which is modified to feature an easy-to-use graphical user interface (GUI), automation, and new parameters for scaling, optimizing the print and projection, and so on. The software is responsible for creating slices of a 3D object given in the form of a STL file and then projecting those slices onto the vat. Firstly, the given STL file is voxelized with consideration of parameters such as the kinematics of the resin, projector specifications, and others (45). This is followed by optimizing and initializing the file using concepts of Radon and inverse Radon transform, biaxial thresholding, Fourier slice theorem, fast-Fourier transform, and more (45). The final step is the generation of a set of 2D images saved as .mat files, which are then continuously looped through in the projection process.

### Acoustic Hardware

The chassis used for the acoustic setup was designed in Solidworks and fabricated using Formlab Form 3+ and Ender 3 S1 3D printers. The glass slides (Sigma Aldrich part number: CLS294775X25) were reshaped using a glass cutter. Various frequency PZTs (Steminc, part numbers: SMPL20W15T14R111, SMPL20W15T21R111, and SMPL20W15T3R111) were first soldered to the connecting wires and then glued alongside the glass slides onto the fabricated chassis. The connections from the PZT were then connected to the function generator (Tektronics TTi AFG3011C) through an amplifier (Digitum electronics, part number: 85184030) to amplify the signal up to 60 V_PP_ at their resonance frequencies.

### Overall Control Unit

The control of the setup is realized by the developed GUI. The projector is connected directly to a laptop/PC via an HDMI cable. An Arduino Mega 2560 with a CNC motor shield and A4988 motor drivers (Conrad, respective part numbers: 191790-AW, 1646889-AW, 1646889-AW) is controlling the motors, and it is also connected to the laptop/PC. Furthermore, the GUI is used to control the function generator for the PZTs control and thus automate and optimize all aspects of the process chain.

### Resin composition and preparation

Both di-pentaerythritol pentaacrylate and aliphatic urethane diacrylate were formulated with 0.6 mM phenylbis (2,4,6-trimethylbenzoyl) (Sigma Aldrich respective product numbers: 407283 and 511447). To achieve homogeneous mixing, they were heated to 100 degrees and simultaneously magnetically stirred at 500 rpm for up to an hour (45, 57).

### Microparticles incorporation and its mixing

The initial experiments used 5-50 micron glass beads (Cospheric part number: SLGMS-2.5 5-50µm - 10g), 15-micron polystyrene beads (Polysciences Polybead microsphere 15 µm catalogue number 18328-5), 1-22 micron stainless steel microspheres (Cospheric part number: SSMMS-7.8 1-22um - 10g), 92 to 110 micron stainless steel microspheres (Cospheric part number: SSMMS-7.8 92-110um - 5g), 90-106-micron fluorescent glass beads (Cospheric part number: HCMS-SLGMS-FMB 90-106um), 106-122 micron glass beads (Cospheric part number: SLGMS-2.5 106-122um - 10g), and 250-300 microns glass beads (Cospheric part number: SLGMS-2.5 250-300um - 10g). We created homogeneous mixtures by using magnetic stirrers at 80 °C and 500 rpm to distribute particles with different concentrations of 20-60 micrograms of microparticles in 20 grams of resin in order to achieve patterns of variable thicknesses in the resin. Before patterning, the ink was centrifuged for 15 seconds at 500 rpm and kept still for up to 10 mins in order to eliminate bubbles and lower the resin temperature. Subsequently, resin with microparticles in the vat was placed in the acoustic setup under volumetric printing conditions and observed under the Zeiss Inverted Microscope (Axiovert 200M) and Leica Inverted Microscope (DMI6000 B) equipped with cameras (Carl Zeiss AxioCAM MrM, and Hamamatsu C11440) for the characterization of the obtained acoustic patterning.

### Data preprocessing

Solidworks was used to create the CAD models that were used as initial input for the volumetric software. Mario, Pharaoh, Nefertiti, liver, lungs and heart geometries were obtained from Thingiverse(58). These files were given to the software in the form of STL. Some of the DiY SonoPrint hardware parts, such as the mechanical fixtures, were designed in Solidworks and sliced using CURA and PreForm software’s before 3D printing them with an Ender 3 S1 and Form 3+ printers, respectively.

### Calibration procedure for SonoPrint printer

Acoustic setups are calibrated using an oscilloscope (Tektronix TBS 2000 series digital oscilloscope 70 MHz 1 GS/s) which is used to measure the resonance frequency of PZT. In addition, for the calibration of the rotational mechanism, we employed high-precision fabrication of the components and calibrated it with a leveler (Emil-Lux Article number: 575459). It was necessary for us to design and fabricate a focusing system to ensure the light is focused at the center of the rotational mechanism. This enabled us to establish the focusing plane. To calibrate the resin-photo initiator concentration, the absorbency spectrum was calculated on the basis of previous literature (45, 57).

### Post processing procedures

Upon completion of printing, isopropanol was used to clean the print. The cleaning was carried out using vibrometer for 10 seconds repeated five times. After cleaning, the printed structure was placed in vacuum for up to 20 minutes with UV light. This step was carried to minimize the oxygen inhibition at the structures surface. Later the structure was exposed to UV light for up to 30 minutes to post-cure the print in UV chamber. Following that, the print was placed in an oven at 85 °C for 60 minutes to improve the strength of the printed component. Lastly, the print was again exposed to UV light for up to 20 minutes.

### Pattern Characterization

After printing, the parts were examined under a microscope with cameras. We also analyzed the fabricated geometry with SEM (used from facility at BRNC, ETH Zurich). Images taken with microscopes are processed using ImageJ software. For recording the patterning inside the printed geometry, cameras from Blackmagic Design and Canon were used. Test methods included tensile using a Zwick 1474 testing machine at 2.5 kN force and compression using a Zwick 1475 testing machine at 100 kN force conducted at RMS Institute, Bettlach. The samples were fabricated according to ASTM standards, namely DIN 604 - (a.o. DIN EN ISO 3167) for compression testing and DIN EN ISO 6892-1 for tensile testing.

## Acknowledgments

We thank Patrick Cologon, Christophe Desarzens, Sarthak Mitter, Tomas Garo Gato, and Pascal Graber for their help in optimizing the hardware and software of the SonoPrint printer. We also thank Zhiyuan Zhang, Mahmoud Medany, and Dimitar Boev for their helpful discussions on acoustic and volumetric setups. We also thank Designing Alley Company for helping us to create schematics.

## Author Contributions

D.A. conceived and supervised the project. P.A., S.Z., S.D., and S.M. performed all the experiments with feedback from D.A. P.A. performed the data analysis, designed, and built the system with feedback from D.A. P.A. and D.A. wrote the manuscript. P.A. drew all the figures. All authors contributed to the scientific presentation and discussion and reviewed the manuscript.

## Competing Interest Statement

The authors declare no competing interests.

## Data Availability

The information supporting the findings of this study can be found within the paper and the Supplementary Information. Further supporting data generated during this study can be obtained from the corresponding author upon reasonable request. The Mario, and Nefertiti model geometries were obtained from Thingverse (under the CC BY 4.0 license).

## Funding

This project has received funding from the Swiss National Science Foundation (SNSF) under the SNSF Project funding MINT 2022 grant agreement No 213058, the European Research Council (ERC) under the European Union’s Horizon 2020 research and innovation program grant agreement No 853309 (SONOBOTS) and ETH Research Grant, grant agreement No ETH-08 20-1.

## Supporting Information for

### Supporting Information Text

#### Supporting Text 1: Theoretical analysis of acoustic setups

Here, we calculate the acoustic wave efficiency for the two piezoelectric transducer (PZT) acoustic setup. A two-dimensional side view of the two PZT setup is shown in **Fig. S1**, where *d* denotes the distance between PZT and the glass resin container, *d_0_* the thickness of the glass resin container and the radius of the glass resin container is denoted by *D/2*. In the transversal plane, a planar two-dimensional rectangular coordinate system is established as seen in **Fig. S1*A***.

In our case for two PZT setup, we have *d* = 2 mm, *D* = 23 mm, *d*_0_ = 0.5 mm.

**Figure S1.**
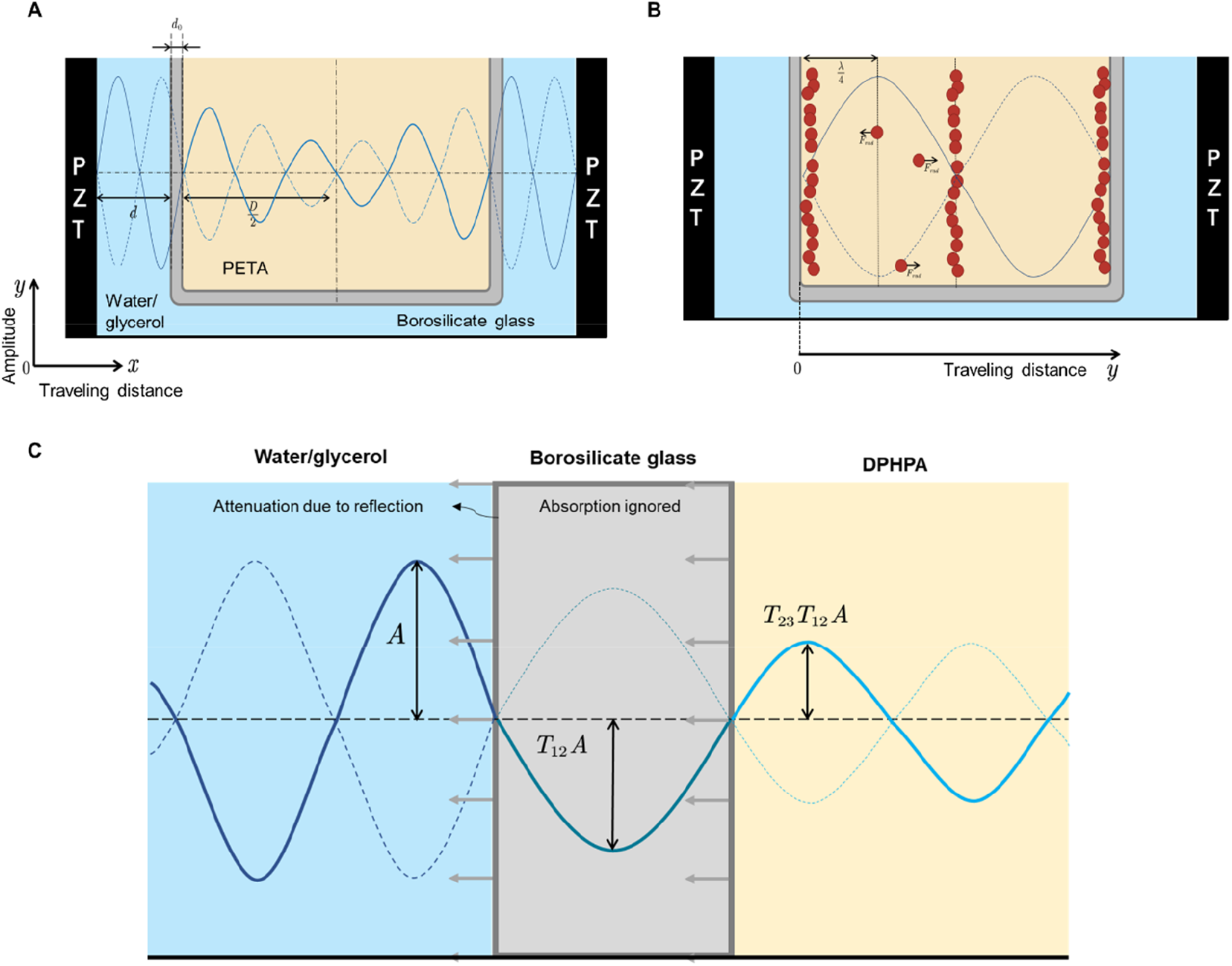
Two-dimensional side view of the two PZT setup. **A.** Schematics showing the standing waves. **B.** Illustration presenting the acoustic radiation forces on microparticles that move them inside the setup when standing wave is applied. **C.** Two-dimensional representation of acoustic wave propagation in the acoustic setup.

### 1.1 Without acoustic attenuation

In the two PZT setup, a standing wave is formed by the superposition of traveling waves from both PZTs. By, considering the traveling wave equation, the standing wave equation is given as (1):

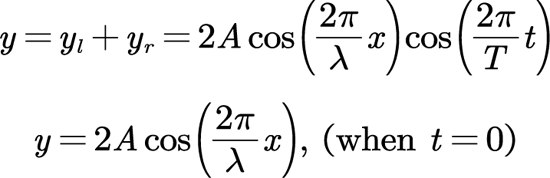

Since 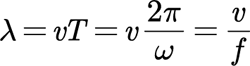
, where v is the acoustic wave velocity inside different media, f is the PZT excitation frequency. We can easily obtain the acoustic wave equation in the acoustic setup:

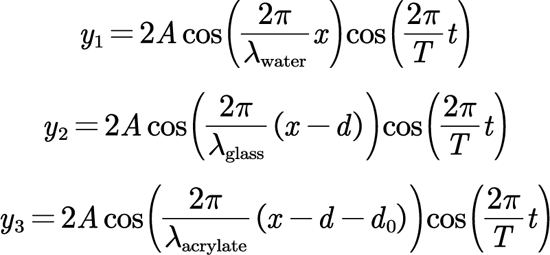

### 1.2 With acoustic attenuation

Acoustic attenuation in a Newtonian fluid follows the Stokes’ law. According to Strokes’ law the amplitude of a plane wave decreases exponentially with distance traveled by the wave, at a rate α which is given by:

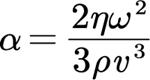

where *η* is the dynamic viscosity coefficient of the fluid, *ω* is the acoustic’s angular frequency, *ρ* is the fluid density, and *v* is the speed of acoustic in the medium.

Without the attenuation effect, the acoustic wave amplitude which is constant throughout the setup is denoted as *A*. When attenuation effect is considered, the amplitude keeps decreasing as the acoustic wave propagates through the water/glycerol medium, glass medium and then through the di-pentaerythritol pentaacrylate resin (PETA) medium.

Stokes’ law cannot be implemented when acoustic wave propagates from water/glycerol to glass, and from glass to PETA mediums. Also, when the acoustic wave meets the intersection of glass from glycerol medium, a part of it is reflected, a part passes through, and he rest is transferred to the glass. As glass only absorbs acoustic energy near its resonant frequency (1) (∼1.837×1012 Hz) (2) as the due to the thinness of the glass resin container, the attenuation inside the glass can also be ignored. Therefore, in our case (1MHz) we only consider the transmission coefficient of glass with respect to acoustic wave.

Assuming that the acoustic wave propagates vertically through the glass vial. The reflection and transmission coefficient of amplitude can be defined by (3, 4):

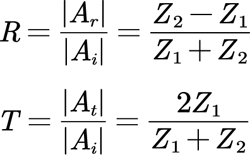

where *A_r_*, *A_t_, A_i_* are the amplitude of the reflected, transmitted and incident wave respectively and, *Z*_1_= *ρ*_1_*v*_1_, *Z*_2_=*ρ*_2_*v*_2_ are the acoustic impedance of the first medium and the second medium. *ρ* and *v* denotes the medium density and acoustic velocity inside the medium (4).

A similar principle is applied, when acoustic wave travels from glass to acrylate for the transmission coefficient. The acoustic wave amplitude change through different medias in this case is thus defined by (3):

- The water/glycerol medium: *A_1_(x) = Ae^−α,x^*, where 0 ≤ *x* ≤ *d*
- The glass medium: *A_2_(x) = T_12_A_1_(d)*, where *d ≤ x ≤ d + d_0_*
- The acrylate medium:

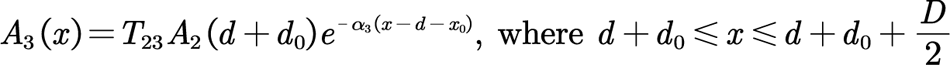

where α1, α3 denote the rate parameter inside water and PETA medium respectively. Moreover, at different temperatures, the change in amplitude with respect to traveling distance is shown in Fig. S2A.

The acoustic wave equation is denoted by:

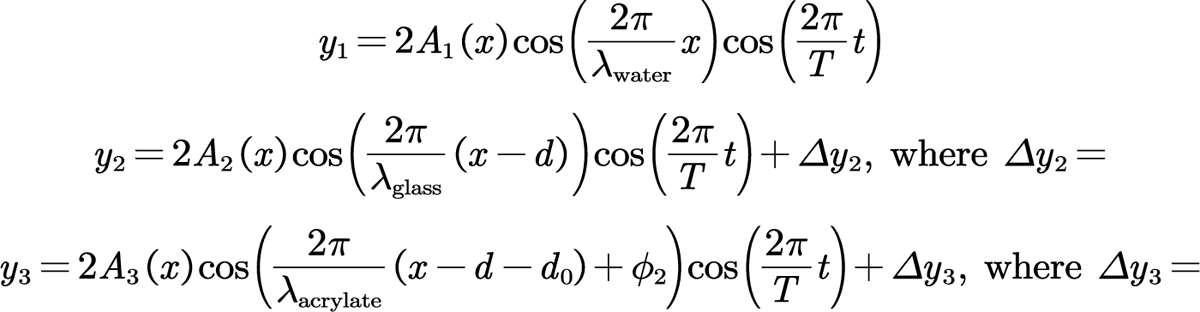

Figs. S2*B* and *C* shows the graphs of acoustic wave attenuation in PETA and glycerol respectively in accordance to distance travelled. In addition, large amplitudes of the transmitted wave are sometimes counterintuitive. However, the energy transported in an acoustic wave is given by (4, 5):

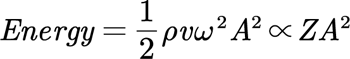

Even with an enhanced amplitude of a transmitted wave generated in certain situations, the energy is conserved.

The ratio of incoming to reflected energy is denoted by *E_R_* and the ratio of incoming to transmitted energy is denoted by *E_T_*. **Figs. 2*D*** and ***E*** depicts the acoustic wave curve with acoustic attenuation effect, acoustic energy attenuation when observed over traveled distance respectively. With respect to impedances on either side of the interface, the energy ratios are denoted by (5):

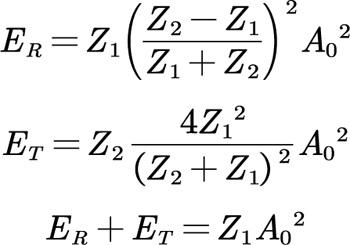

**Figure S2.**
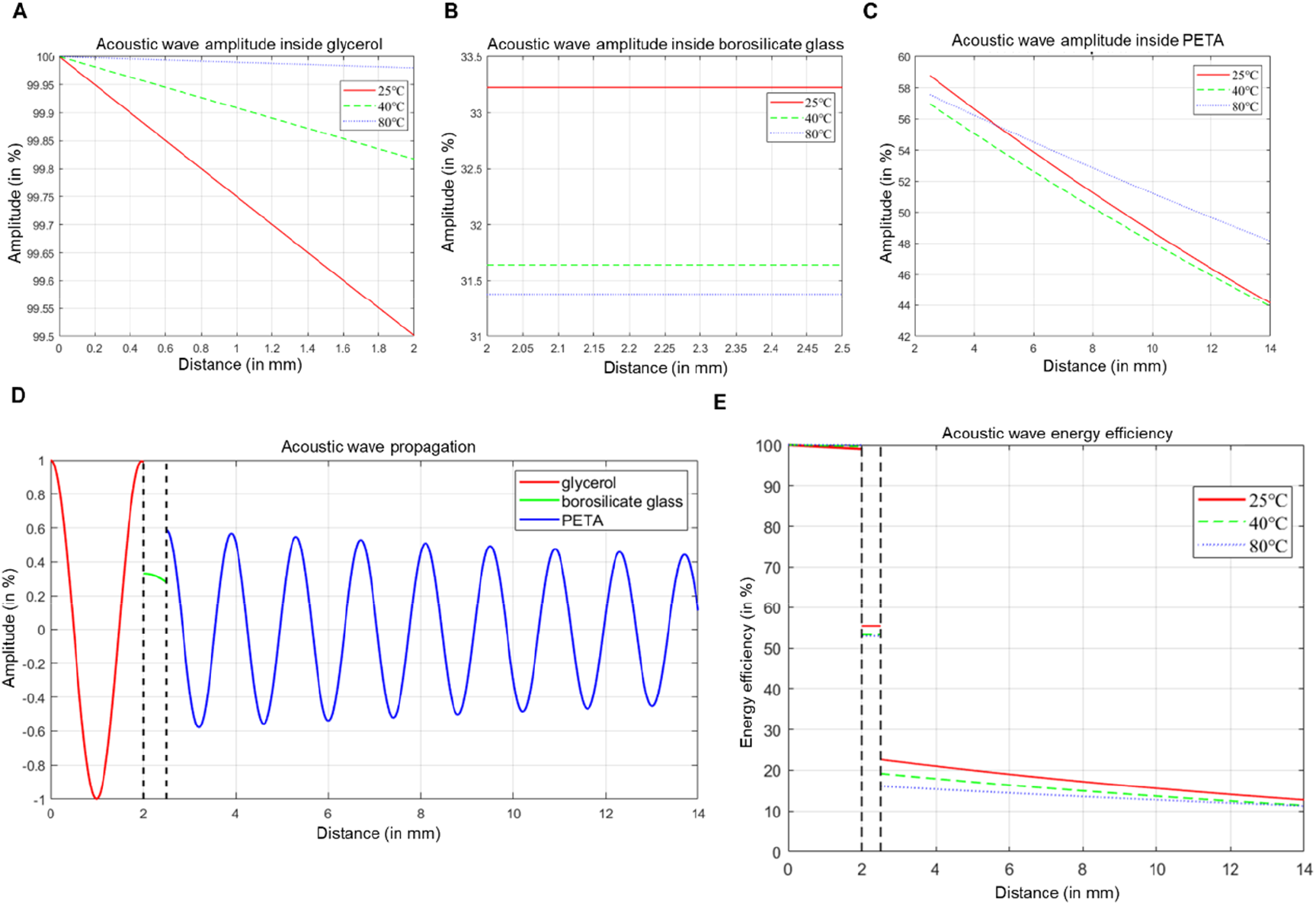
Graphs with acoustic wave amplitude vs distance travelled in the two PZT acoustic setup. **A, B** and **C**. Graph showing the attenuation of acoustic wave amplitude with distance traveled considering the two PZT acoustic setup in glycerol, borosilicate glass and di-pentaerythritol pentaacrylate resin (PETA) respectively. D. Shows the graph with acoustic wave amplitude in % vs distance travelled. E. Graph depicts the acoustic energy efficiency in % vs distance traveled.

### 1.3 Acoustic patterning of microparticles

#### 1.3.1 Working principle

The two PZT acoustic setup consists of two PZTs in a parallel arrangement. A solution of microparticles is infused into a cylindrical vial placed in the center, surrounded with a water/glycerol medium. The two PZTs generate two series of acoustic waves, whose interference generate a stable standing wave, with a periodic distribution of pressure nodes and anti-nodes (6).

### 2.2 Quantitative force analysis

Microparticles in fluid medium when exposed to the acoustic standing wavefield are subjected to acoustic radiation force (ARF) and acoustic drag force. The radiation force pushes the microparticles to move, where as, acoustic drag force, acts opposite to the relative motion of the moving microparticles. Under acoustic standing wave field, the particles will move to the either to the nearest node or antinode depending upon the contrast factor.

The primary ARF with respect to acoustic wave traveling distance (y) acting on a compressible, spherical suspended particle in an arbitrary acoustic field is given by Gor’kov’s theory and simplified by Yosioka’s equation when the radius of the particle a is much less than the wave length λ (7):

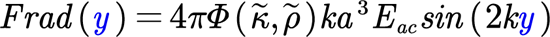

where

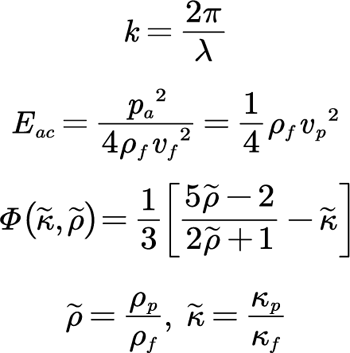

*E_ac_* is the acoustic energy density, *Φ*(*κ*°*, ρ*°) is the acoustic contrast factor, *k* is the wave number, *ρ_p_*, *ρ_f_* and *κ_p_*, *κ_f_* represent the density and compressibility of the particle and fluid respectively, and *v_p_* is the velocity of the particle inside the fluid.

The acoustic contrast factor dictates the minimum position of the Gor’kov potential (7). If its value is positive, the microparticle will be pushed towards the pressure node of the wave field and vice versa (8). At room temperature of 25, the density of PETA is 1155*kg/m*^3^. The commercially available stainless steel metal microspheres with a diameter of 1-22 *µm* has a density of 7700-7900*kg/m*^3^ (available from the technical data sheet from Cospheric (9)). *κ*° can be obtained by the equation

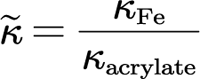

The compressibility of metal is 5.88×10^−7^ Pa^−1^ (10). The compressibility of PETA, which equals the inverse of the product between density and acoustic velocity, i.e., is estimated to be ∼3.5×10^−10^ Pa−1. Therefore, the acoustic contrast factor of stainless-steel metal microspheres in PETA is positive. Stating that the microparticles will be pushed towards the pressure nodes of the standing wave.

Acoustic drag force is the frictional force exerted on spherical objects with very small Reynolds numbers in a viscous fluid. It is given by Stokes’ drag:

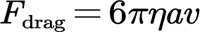

where *η* is the dynamic viscosity of the fluid, *a* is the radius of the spherical particle, *v* is the flow velocity relative to the particle.

In order to determine the time needed to pattern the microparticles, as determined previously we know the metal microspheres will move to nodal position. Considering the particle is located at (**Fig. S1*B***), moving towards the co-ordinate at *y* = 0. In this case, the displacement of the particle is negative as the positive direction of the *y* axis is considered as the positive direction of the system, i.e. 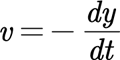.

When the microparticles maintain a constant velocity inside the medium, the acoustic radiation force and the acoustic drag force balance each other, i.e. (6)

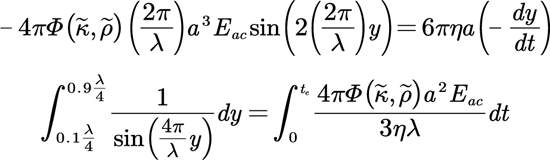

Therefore, the time needed for patterning is given by

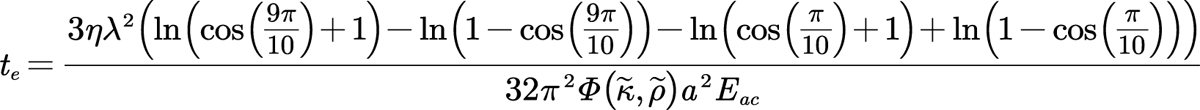

#### Supporting Text 2: Characterization of acoustic patterning

A control experiment was conducted to ensure that acoustic patterning was intact for the entire duration of volumetric printing. For this we used a two PZT setup with PETA and 5-50 µm glass microspheres of resin temperature range up to 40 °C. In order to mimic the exact same conditions as when volumetric printing, we first acoustically patterned microparticles. We then recorded the acoustic patterning using Black Magic camera from a side view for up to ten minutes. Using ImageJ software, we compared the patterning over various periods of time. We observed that the acoustic patterning remained intact throughout the control experiments (see **Fig. S3**). In our setup, the volumetric printing process usually lasts for up to 240 seconds, which indicates that the acoustic patterns remain stable throughout the process.

**Figure S3.**
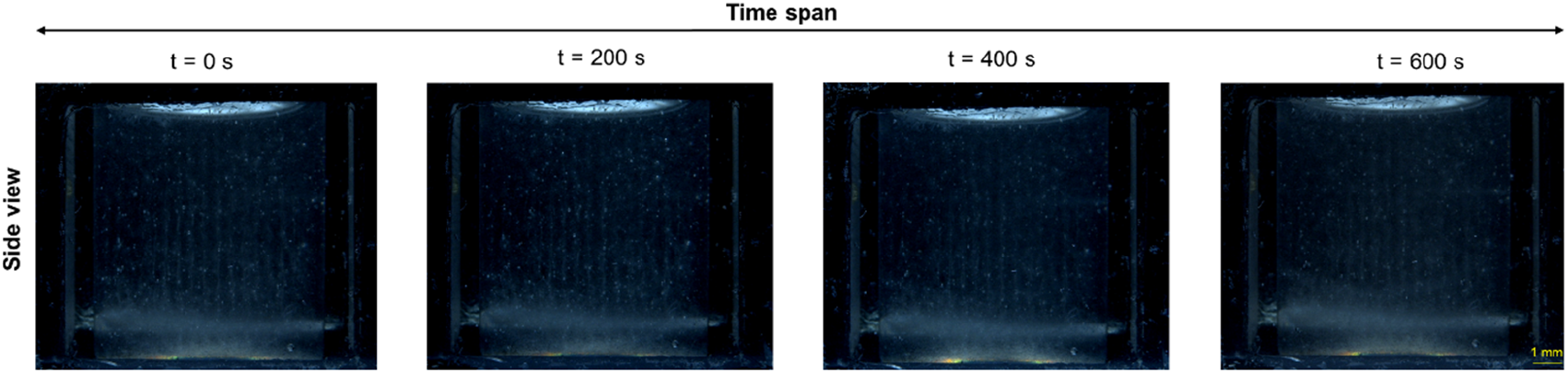
Control experiments to determine the stability of acoustic patterning of microparticles. The images show side view of two PZT setup taken over a time span of 600 seconds. For this experiment we used 1 MHz two PZT setup with 5-50 µm glass microspheres excited at 1.1 MHz, 30 V_PP_, and 40°C. Scale bar: 1 mm.

### 2.1 Two PZT acoustic setup

As a result of changing the PZTs in a setup, acoustic patterning with varying distance between formed patterns could be achieved, as shown in Figs. S4A, B, and ***C***. We used two PZT setup to demonstrate our ability to manipulate the distance between the formed acoustic patterns. As a demonstration, we patterned lines using soda-lime glass microspheres with a diameter of between 5 and 50 µm in PETA. As a result of changing the frequency of the setups, we were able to alter the distance between the formed pattern from 0.96 to 0.45 mm, and the distance can be changed further if a different set of PZTs is used. In **Figs. S4*D*** and ***E***, the top, side and bottom view of the setups are shown also see **Movies S1, S2** and **S5**.

**Figure S4.**
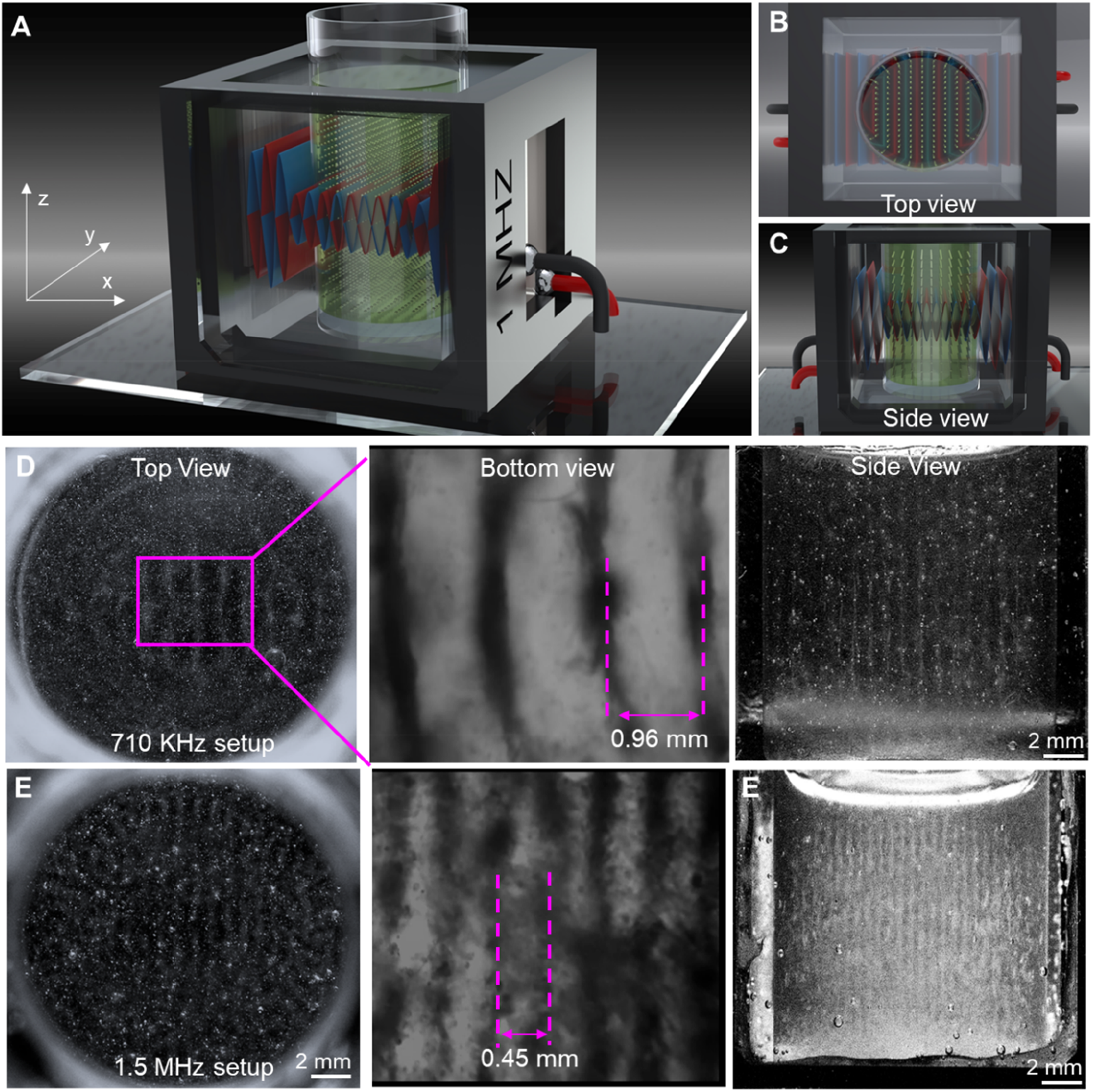
Different resonant frequency piezoelectric transducers (PZTs) for two PZT acoustic setup and the microparticle patterning formed. **A, B,** and **C.** Schematic illustration of two PZT setup showing the bulk acoustic waves and microparticle patterning from isometric, top and side views. **D** and **E.** The images show the top, side and microscopic views of the two PZT setup with 1 MHz and 1.5 MHz resonant PZT creating line formation. Experiments used 5-50 µm glass microspheres mixed in resin. Scale bar: 2 mm for side and top view.

#### 2.1.1 Varied patterns with two PZT acoustic setup

We performed experiments to show that various patterns can be generated by changing the temperature of the resin and by exposing the vial to rapid temperature shock. We observed that microparticles patterned at different temperatures, frequency and voltages results in varied patterns. For example, when a pair of parallel PZTs were excited at 1.02 MHz and 45 V_PP_ at 35°C, they produced a radial pattern as seen from top and side view in **Fig. S5*A***. Also, when the resin temperature was 50°C and the PZTs were excited at 1.03 MHz and 60 V_PP_, a line pattern was produced. Following this, the vial was placed in a cold bath, which was set at a temperature of 4 to 14°C. In response to the sudden change in temperature, the particles began moving towards the center of the vial, when stabilized, resulted in an inwards pattern, as seen from and side view in **Fig. S5*B***. When very high amplitude is applied we were able to achieve a swirling motion as seen **Fig. S5*C***. Here we used 1.02 MHz and 60 V_PP_ at a temperature of 60°C. Lastly, to characterize the rapid cooling with two PZT setup, we formed various line patterns and with resin temperature of 50 to 60°C using the 1.01 MHz and 60 V_PP_ excitation, followed by immediate rapid cooling in a cold bath maintained at 4 to 14°C, resulting in radial line patterning as seen in **Fig. S5*D*** *− **F***.

**Figure S5:**
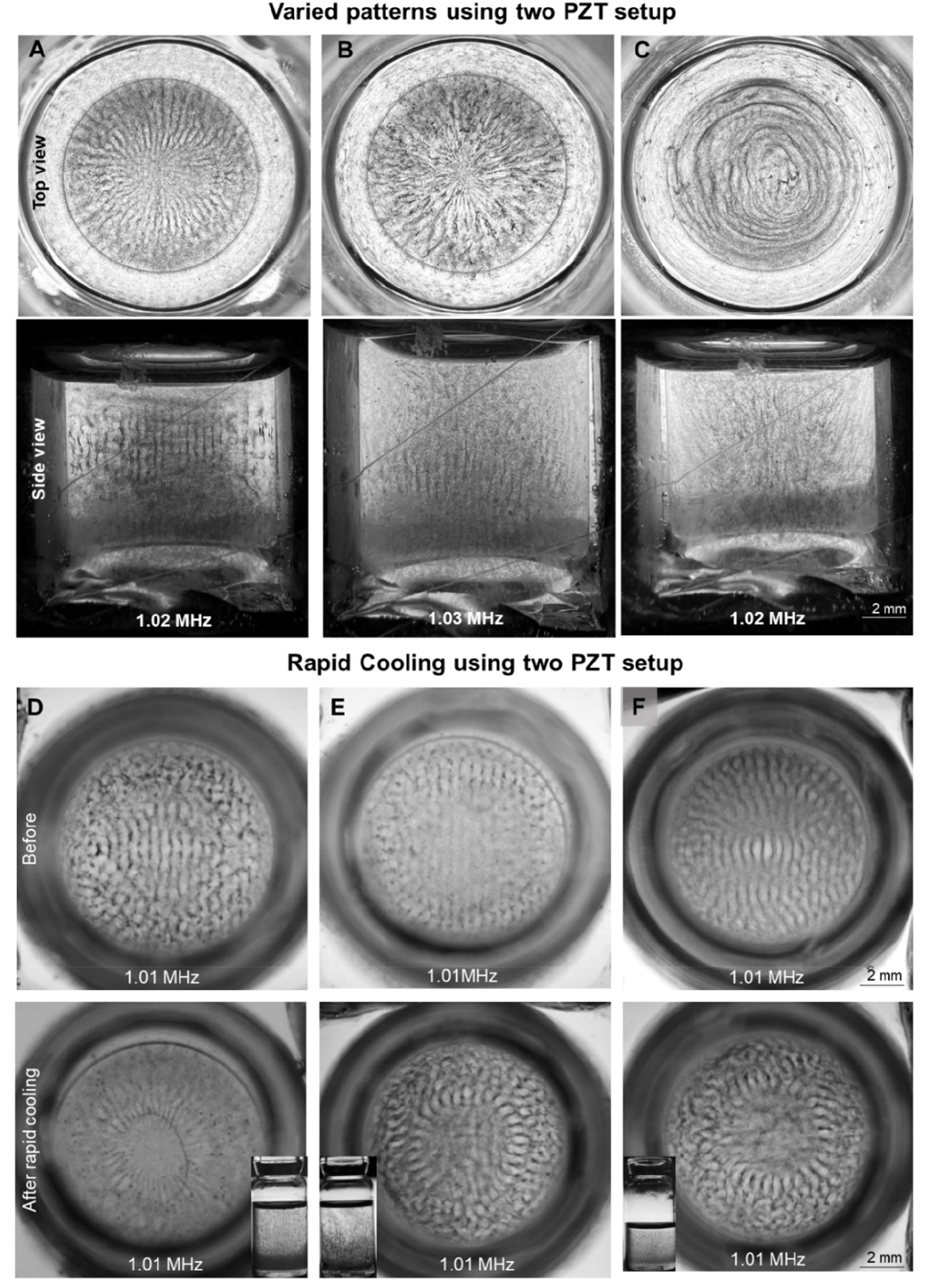
Microparticle patterning using two PZT acoustic setup. **A.** The image shows the top and side view of the 1 MHz two PZT acoustic setup when excited at 1.02 MHz at 45 V_PP_ at a temperature of 35°C. Scale bar: 2 mm. **B.** A top and side view of the pattern generated by two PZT 1MHz setup excited at 1.03 MHz, 60 V_PP_, and 50°C, immediately followed by rapid cooling at 6°C is shown in the image. Scale bar: 2 mm. **C.** Top and side view of pattern generated at 60°C by excited two PZT setup at 1.01 and 60 V_PP_ generating a swirling motion. Scale bar: 2 mm. **D, E, and F.** Shows patterning of microparticles at 50 to 60°C at 1.01 and 60 V_PP_ and then immediate rapid cooling in cold bath maintained at 4 to 14°C, leading to radial line patterning. Scale bar: 2 mm.

### 2.2 Four PZT acoustic setup

The four PZT arrangement incorporates four orthogonally-positioned PZTs; activating these PZTs at a similar frequency produced rhombus patterns in the xy-plane, which were projected along the z-axis as seen in Figs. S6A, B, and ***C***. With this acoustic setup, we were able to produce various patterns through the combination of different pairs of parallel PZTs excited at the same frequency, resulting in three combinations (two parallel PZTs excited at the same time or all four PZTs excited at the same time). Also, we were able to achieve different patterns by using a single pair of transducers that operate at different frequencies, as in the two PZT setup, see **Movie S4**. Furthermore, we were able to obtain different patterns by stimulating all four transducers at the same frequency and then changing the frequency (as shown in **Fig. S6** and **Movie S3**).

**Figure S6.**
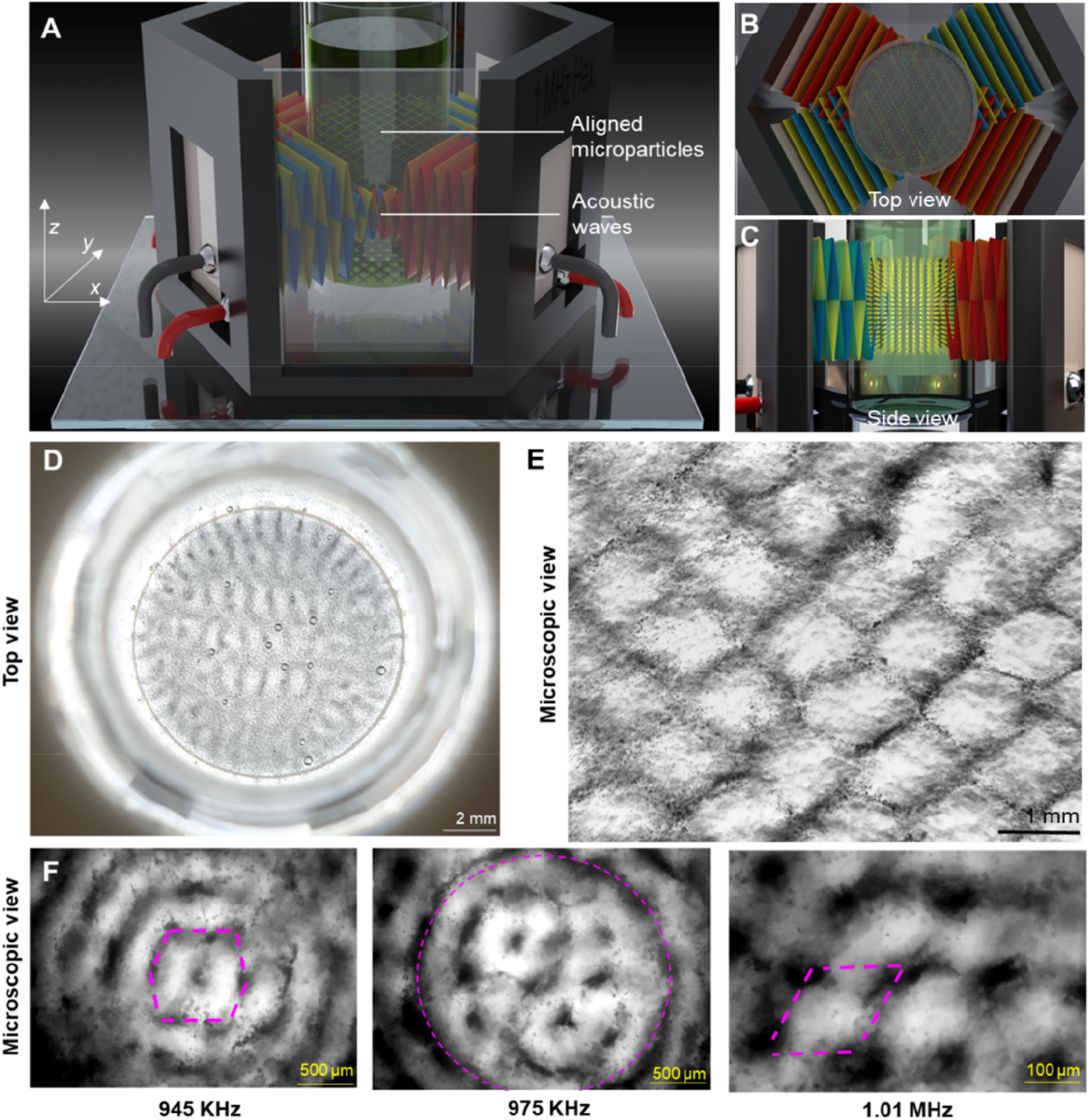
Acoustic patterning of microparticles in four PZT setup. **A.** A schematic representation of a four PZT acoustic setup that utilizes four identical piezoelectric transducers (PZT)s orthogonally positioned in yz-plane being actuated simultaneously at resonance frequency to produce bulk acoustic waves for patterning microparticles in the desired manner. **B.** and **C.** Displays the top and side views of the four PZT acoustic setup shown in A, with microparticles patterned in the form of quadrilaterals along xy-plane and distributed overall in xz-plane respectively. **D**. The images show the top view of the 1 MHz four PZT setup excited at 1.01 MHz at 30 V_PP_. Scale bar: 2 mm. **E.** The figure illustrates the pattern generated by 1 MHz four PZT setup from the microscope view of the pattern formed in D; 15 µm polystyrene particles in PETA were used; also see **Movie S5**. Scale bar: 1 mm**. F**. Displays the changes in patterning of microparticles at 945, 975, and 1010 kHz from the microscopic view with PETA and 5-50 µm glass microspheres excited at 1.1 MHz at 45 V_PP_. Scale bar: 500 µm for left and middle image and 100 µm for right images.

### 2.3 Six PZT acoustic setup

This study utilized a six PZT setup to achieve diverse microparticle patterns. At first, the glass resin vat was positioned within the six PZT setup, and when the PZTs were activated, bulk acoustic waves were generated to precisely pattern the microparticles according to the desired configuration, as shown in **Fig. S7*A***. Following this, the vial containing the aligned microparticles was transferred to a volumetric printing setup as shown in **Fig. S7*B***. The printing arrangement included an index-matching container made of glass with a refractive index of 1.516, filled with glycerol serving as the index-matching fluid. A control experiment was conducted to characterize the change in patterning based on changes in distances, frequencies and activation of PZTS on the formed patterns, see **Fig. S7*C-F***.

**Figure S7.**
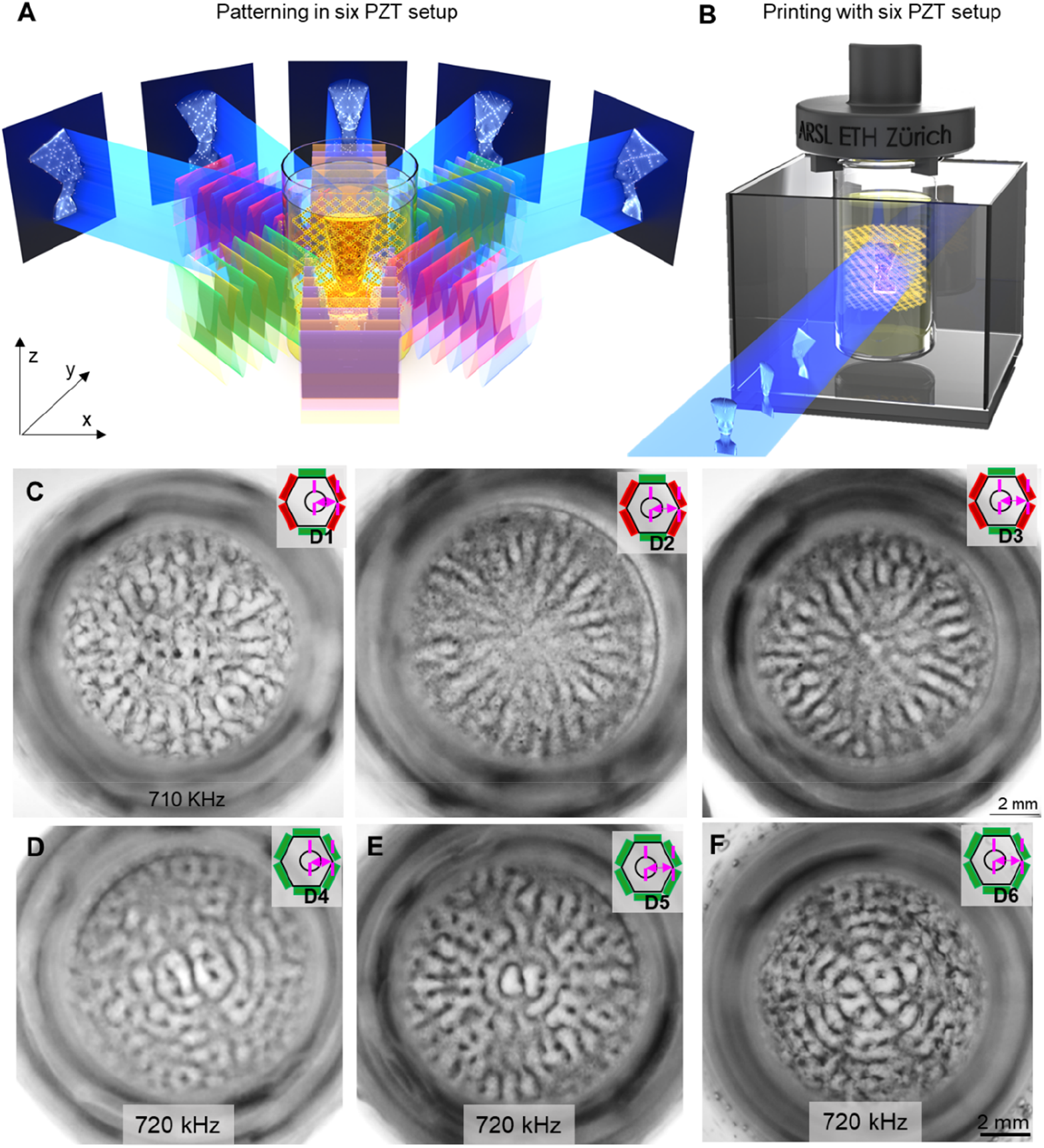
Acoustic patterning using six PZT setup. **A.** Schematics shows patterning of microparticles when all six PZTs are activated in the six PZT setup. It also shows the images generated and projected at different angles for fabricating a Nefertiti model geometry. **B.** Schematic depicts the transfer of glass resin vat container after acoustic patterning to a volumetric compatible chamber for volumetric printing of a Nefertiti model geometry. **C.** Shows the top view images when the setup is excited at 710 kHz and the changes in vial position from D1 – D3 where it corresponds to ∼21.6, 18.48, 19.32 mm respectively. Scale bar: 2mm. **D – F.** Shows the top view when all six PZTs were excited at 720 kHz, 30 V_PP_. Where D4 – D6 corresponds to ∼18.9, 21.8 and 22.7 mm respectively. Scale bar 2 mm.

#### 2.3.1 Temperature shock in six PZT acoustic setup

In order to evaluate the pattern formation caused by temperature shock, we conducted temperature shock experiments on particles without acoustically patterned surfaces. As shown in **Fig. S8**, in the presence of a temperature shock from 60 to 6 °C, the particles do not form a pattern. Furthermore, as we performed for the two PZT acoustic setups, we also conducted temperature characterization experiments for six PZT setups, concluding that upon temperature shock, patterned microparticles move towards the center, forming radial patterns for all different kinds of initial patterns formed as seen in **Fig. S9**.

**Figure S8.**
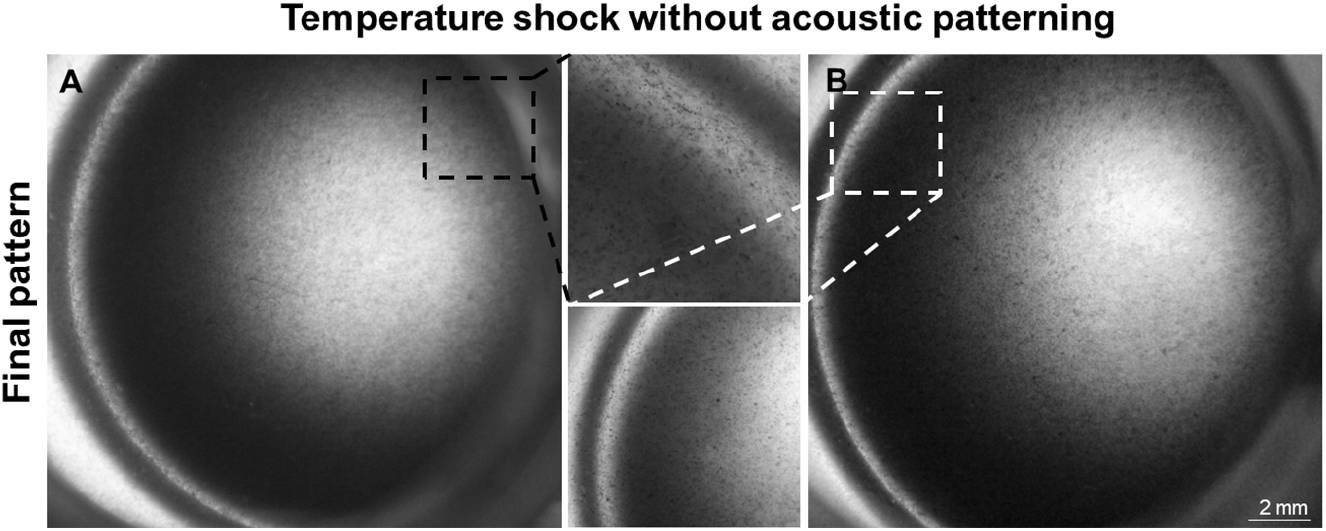
Temperature shock to microparticle without patterning. **A.** and **B.** Shows images of top view of glass resin vat with microparticles after temperature shock from 60°C to 6°C

**Figure S9.**
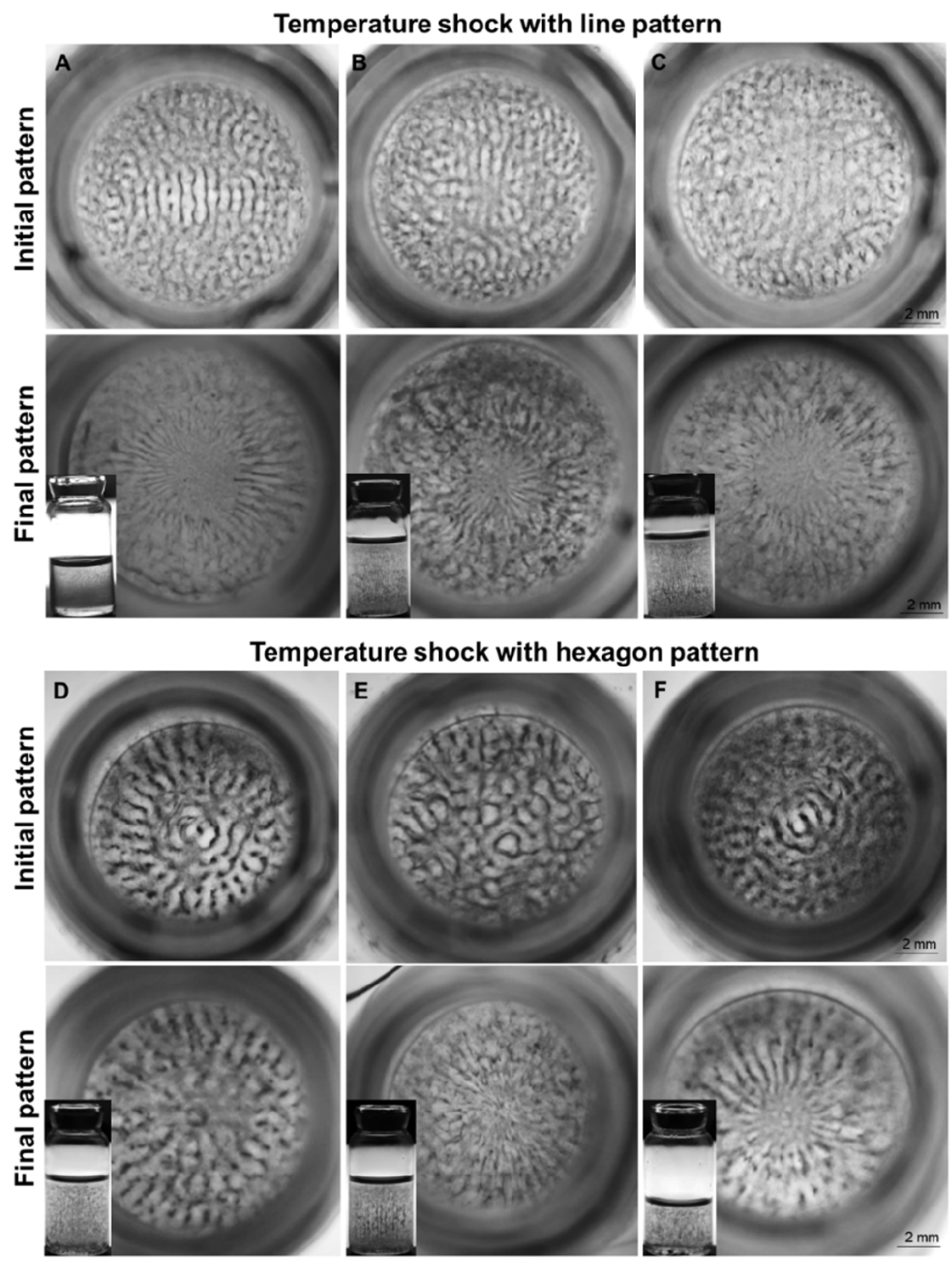
Temperature shock to microparticle patterning formed using six PZT acoustic setup. **A, B** and **C.** Line patterns formed by actuating a pair of parallel PZT at 715 kHz, 60 V_PP_, and 50°C and then a temperature shock was given to 10°C the microparticle move towards the center to form radial line pattern. Scale bar: 2 mm. **D, E** and **F.** Polygonal patterns formed by actuating all six PZTs at 715 kHz, 60 V_PP_, and 50°C and then a temperature shock was given to 10°C the microparticle move towards the center to form radial line pattern. Scale bar: 2 mm.

#### **Supporting Text 3**: Printed geometry characterization

To assess the influence of microparticle concentration on the printed structures, we fabricated cylindrical geometries using PETA resin with varying concentrations of 1-22 microns stainless steel microspheres, as illustrated in **Fig. S10**. These experiments allowed us to gain valuable insights into the interplay between temperature, microparticle concentration, and the overall printing results.

To analyze the consistency of the patterns inside a volumetrically fabricated print we printed a cylindrical geometry to assess the microparticle patterning at different depths and its consistency throughout the 3D structure as shown in schematics of **Fig. S11*A***. We first excited a pair of parallel PZTs at 1.01 MHz and 30 V_PP_ at 25°C to pattern 1-22 µm stainless steel microspheres in PETA, followed by 90 seconds of printing over the patterned microspheres. The printed structure was characterized after post-processing. An analysis of the structure from the side view at different angles shows a sheet of patterned particles separated by a certain distance (see **Fig. S11*B***). When viewed from 90- and 270-degree angles, lines with a spacing of approximately 0.7 mm between consecutive lines can be observed. While at 180- and 360-degrees, microparticles are seen distributed over the entire plane. Under a microscope, the patterning was examined along three distinct depth planes, I, II, and III, respectively, along the printed structure; see **Fig. S11*C***. **Fig. S11*D*** shows the structure’s top view. As a result, patterning consistency was achieved throughout the printed geometry.

Additionally, we demonstrated the capabilities of composite fabrication by showing a few potential applications for the future. Here we first show composite applications for aerospace and automotive industries using 1-22 µm stainless steel metal microparticles in PETA resin patterned using 1MHz two PZT setup. As can be seen in **Fig. S11*E***, we demonstrated by printing helicopter propellers from toy helicopter models. A pair of PZTs were excited at 1.02 MHz and 30 V_PP_ to achieve the helicopter propeller. Under a microscope, we examined the patenting as seen in **Fig. S11*E*** bottom right corner. **Fig. S11*F*** illustrates the application of composite materials to the automobile sector through the printing of wheels for a toy car. The pair of parallel PZTs in two PZT were excited at 1.02 MHz at 30 V_PP_ to achieve the line pattern which was then exposed to temperature shock to achieve radial line pattern, and then the wheel geometry was printed. A subsequent analysis of the patterning at the center of the wheels was conducted under a microscope as seen in **Fig. S11*F***, bottom right corner. Furthermore, composite materials can be used in a variety of other industries, such as snowboards, which require high levels of strength and durability. We demonstrate this with a toy skateboard, as shown in **Fig. S11*G***. For patterning the microparticles, we excited the transducers at 1.02 MHz at 30 V_PP_ and then printed the geometry. In the bottom right corner of the figure, a zoomed-in view is shown. All of these patterns and results confirm the many possibilities that the SonoPrint printer may offer. In addition, it shows its potential to open up a new application for volumetric printing technology in the composite fabrication industry.

**Figure S10.**
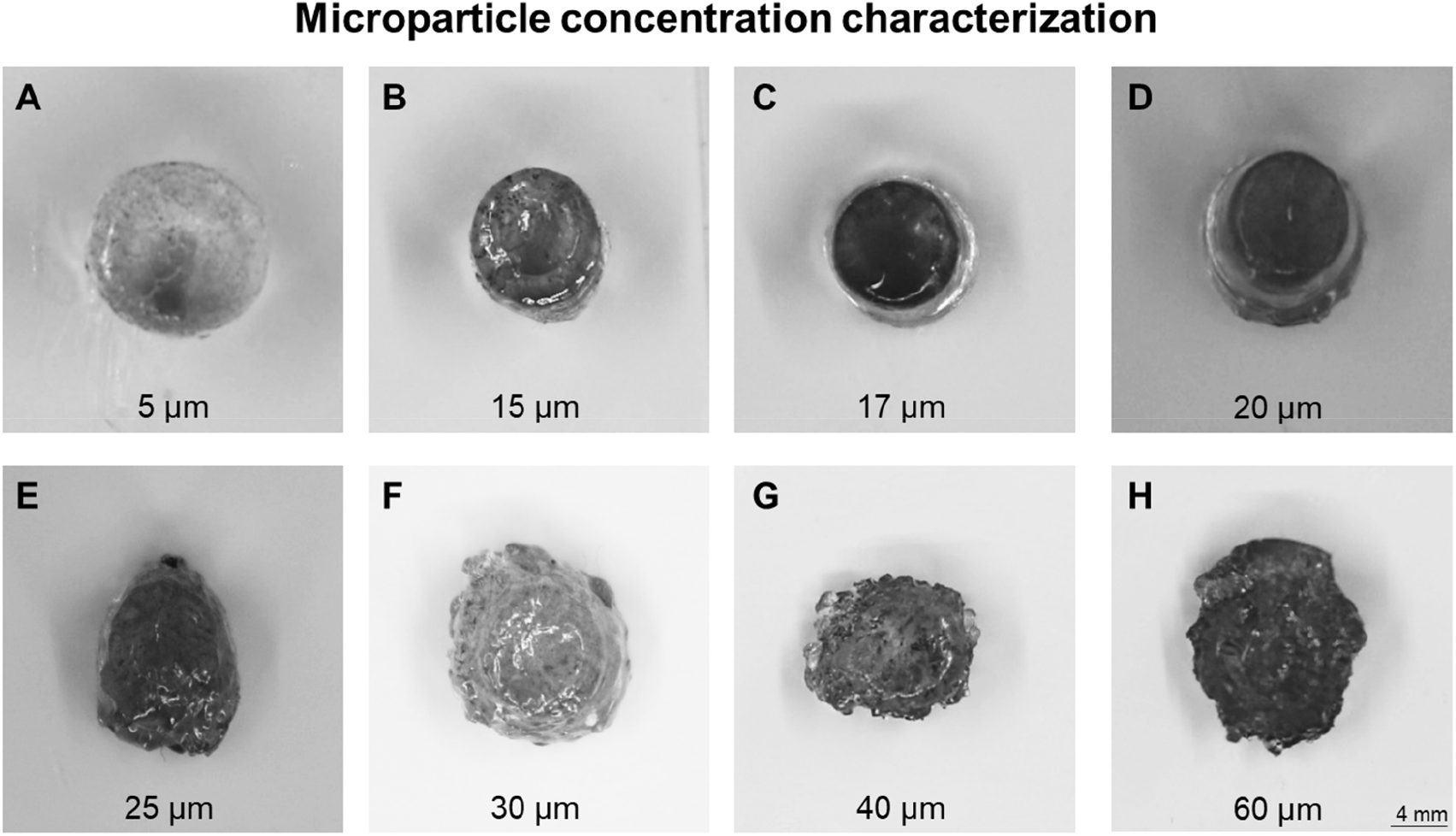
Microparticle concentration characterization for volumetric printed structures. **A-H**. Volumetric printed structures different 1-22 microns stainless steel microsphere concentration in 4 grams of PETA resin. Scale bar: 4 mm. Error bar: 5.

**Figure S11.**
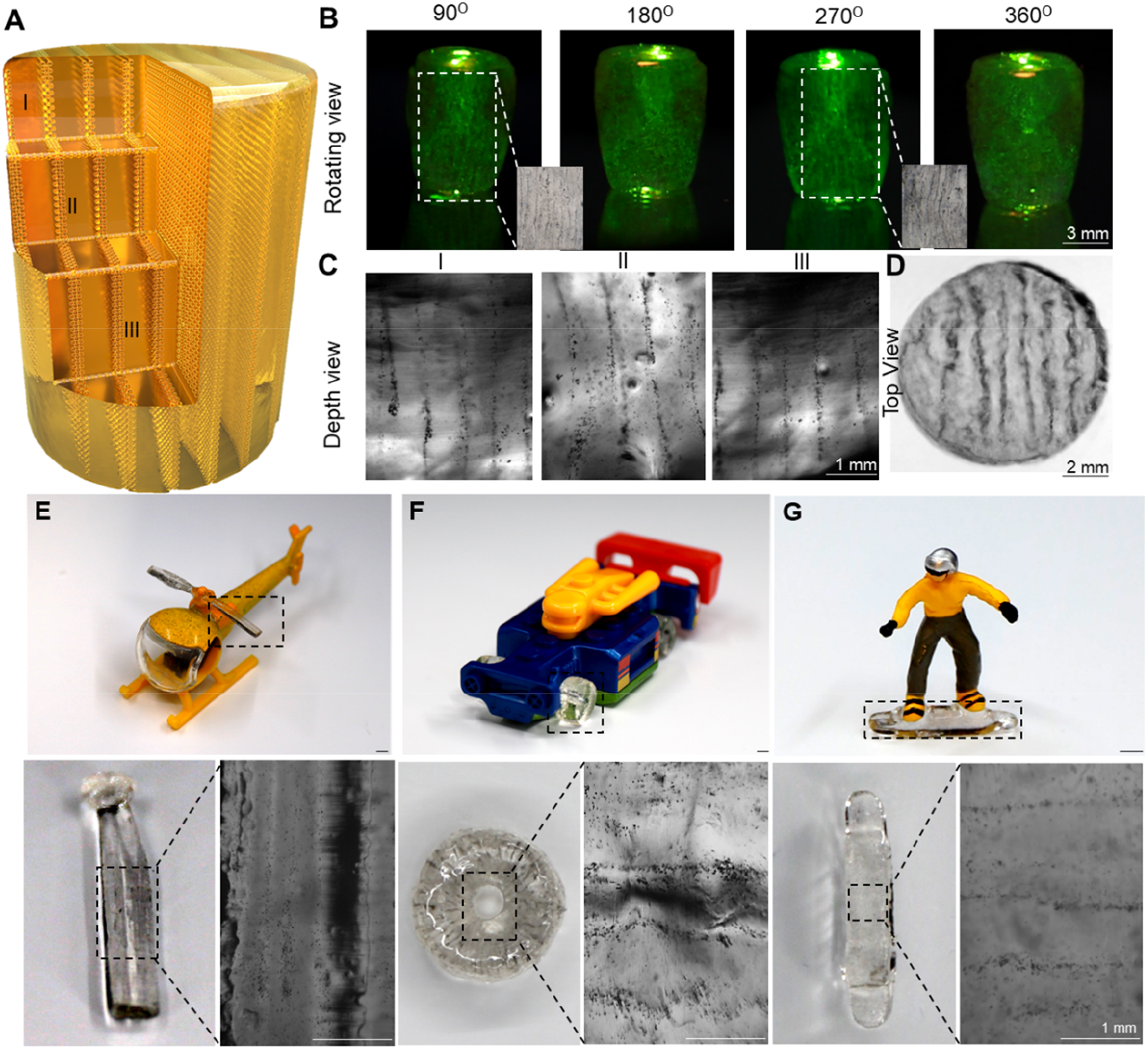
The characterization and applications of SonoPrint printers’ printed structures. **A.** An illustration of the cut section across three different planes I, II, and III of cylindrical model geometry. A pair of piezoelectric transducers (PZT)s were excited at 1.01 MHz and 30 V_PP_ using 1-22 μm stainless steel metal microspheres in PETA. **B.** The images depict the side view of the fabricated cylindrical structure from four different angle perspectives 90^0^, 180^0^, 270^0^, and 360^0^ respectively. Scale bar: 3 mm. **C.** Displays a microscopic view of the cylindrical structure when seen at depth planes I, II and III. Scale bar: 1 mm. **D.** Displays the top surface view of the cylindrical structure. Scale bar: 3 mm. **E.** Demonstration of a helicopter model featuring SonoPrint fabricated main rotor blades. The fabricated main rotor blade is shown in the bottom left corner and its zoomed-in view appears in the bottom right corner. Scale bar: 1 mm. **F.** Demonstration of a car model featuring SonoPrint-fabricated wheels. The bottom left corner shows the fabricated wheel composite geometry, while the bottom right corner shows a zoomed-in view. Scale bar: 1 mm. **G.** The SonoPrint-fabricated skateboard is demonstrated with a man’s model. On the bottom left, the fabricated skateboard composite geometry is displayed, while on the bottom right, a zoomed-in view is shown. Scale bar: 1 mm.

## Supporting Tables

**Table S1:** Parameters for density, viscosity and wave velocity.

| Parameter | Media | 25°C | 40°C | 80°C |
| --- | --- | --- | --- | --- |
| **Density (kg/m<sup>3</sup>)** | water | 997.05 | 992.25 | 971.60 |
|  | glycerol | 1257.7 | 1248.5 | 1224.0 |
|  | PETA | 1155 | ~1150.2 | ~1129.55 |
|  | glass | 2.23e3 | 2.23e3 | 2.23e3 |
| **Dynamic Viscosity (Pa.s)** | water | 8.90e-4 | 6.5272e-4 | 3.5404e-4 |
|  | glycerol | 9.0568e-1 | 2.8244e-1 | 3.1500e-2 |
|  | PETA | ~7.4 | ~7 | ~5 |
|  | glass |  |  |  |
| **Acoustic wave velocity (m/s)** | water | 1474 | 1526 | 1555 |
|  | glycerol | 1964 | 1866.6 | 1885.2 |
|  | PETA | ~1666 | ~1696 | ~1720 |
|  | glass | 5560 | 5560 | 5560 |

**Table S2:**
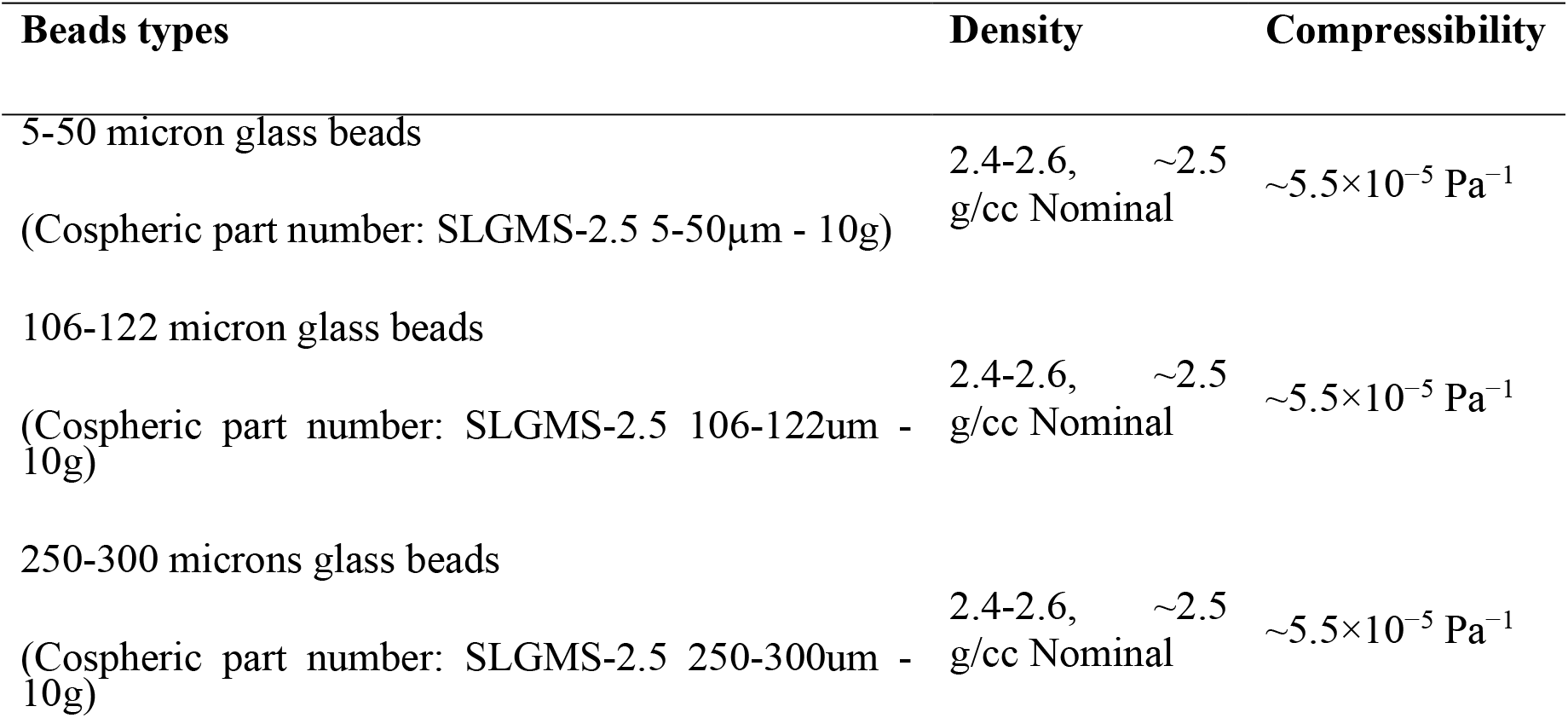
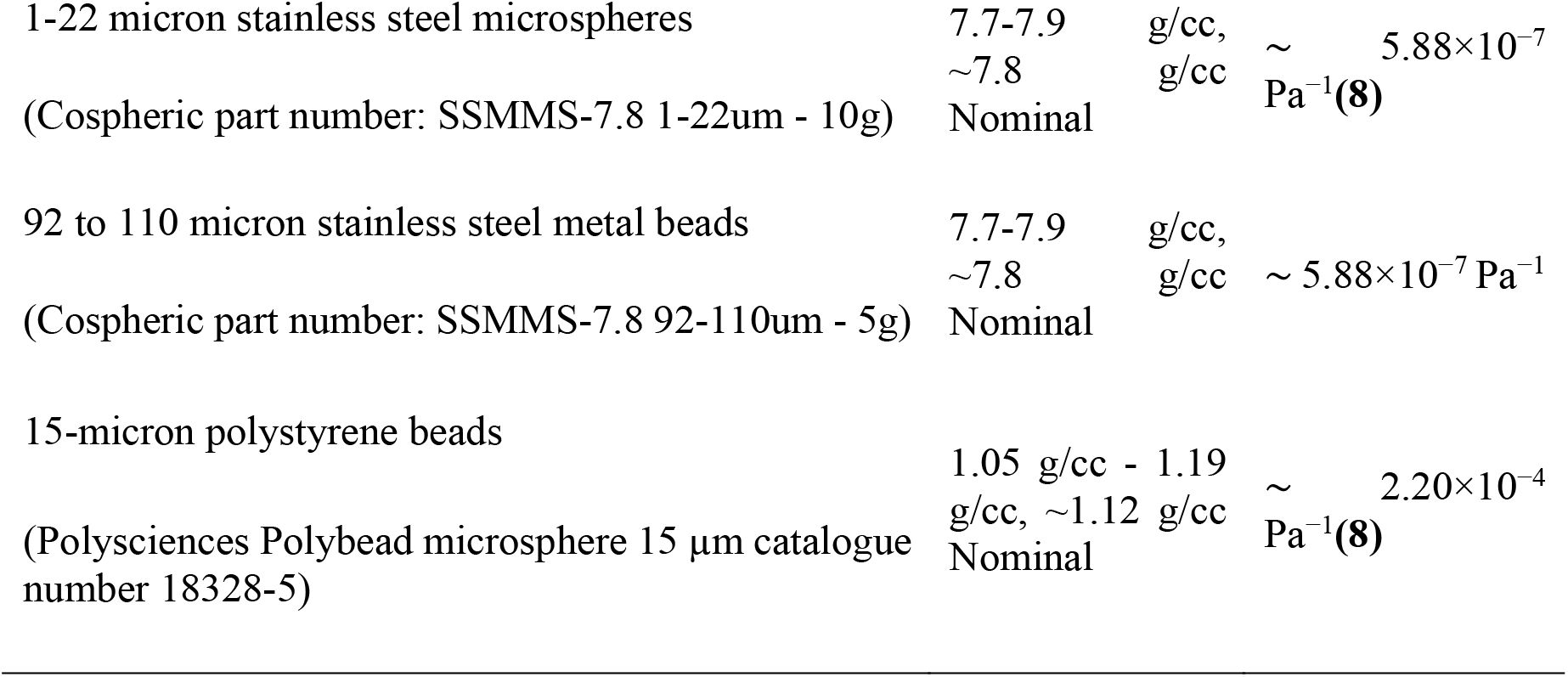
Microparticles parameters.

## Supporting Movie Legends

**Movie S1**: Acoustic Patterning utilizing 1MHz two PZT acoustic setup with 5 – 50 micron soda lime glass microspheres in PETA resin.

**Movie S2**: Top and side view of 1.5 MHz two PZT acoustic setup utilizing 5 *−* 50 micron soda lime glass microspheres in PETA resin.

**Movie S3**: Changing frequency in 1 MHz four PZT acoustic setup using 5 *−* 50 micron soda lime glass microsphere in PETA resin to achieve various patterning.

**Movie S4**: Microscopic view of 1 MHz four PZT acoustic setup utilizing 15 micron polystyrene beads in PETA resin.

**Movie S5**: Changing the concentration of microparticles in 1 MHz two PZT acoustic setup using 5 *−* 50 micron soda lime glass microsphere in PETA resin.

**Movie S6:** Nefertiti composite geometry fabrication using SonoPrint printer.

## Notes

### Competing Interest Statement

The authors have declared no competing interest.

## References

1. R. N. Yancey, “2 - Challenges, opportunities, and perspectives on lightweight composite structures: Aerospace versus automotive” in Lightweight Composite Structures in Transport, J. Njuguna, Ed. (Woodhead Publishing, 2016), pp. 35–52.

2. J. C. Najmon, S. Raeisi, A. Tovar, “2 - Review of additive manufacturing technologies and applications in the aerospace industry” in Additive Manufacturing for the Aerospace Industry, F. Froes, R. Boyer, Eds. (Elsevier, 2019), pp. 7–31.

3. P. D. Mangalgiri, Composite materials for aerospace applications. Bull Mater Sci 22, 657–664 (1999).

4. H. Singh, G. Singh Brar, H. Kumar, V. Aggarwal, A review on metal matrix composite for automobile applications. Materials Today: Proceedings 43, 320–325 (2021).

5. C. E. Bakis, et al., Fiber-Reinforced Polymer Composites for Construction—State-of-the-Art Review. Journal of Composites for Construction 6, 73–87 (2002).

6. I. Paoletti, Mass Customization with Additive Manufacturing: New Perspectives for Multi Performative Building Components in Architecture. Procedia Engineering 180, 1150–1159 (2017).

7. A. Pajonk, A. Prieto, U. Blum, U. Knaack, Multi-material additive manufacturing in architecture and construction: A review. Journal of Building Engineering 45, 103603 (2022).

8. M. S. B. Reddy, D. Ponnamma, R. Choudhary, K. K. Sadasivuni, A Comparative Review of Natural and Synthetic Biopolymer Composite Scaffolds. Polymers 13, 1105 (2021).

9. M. J. Mochane, T. S. Motsoeneng, E. R. Sadiku, T. C. Mokhena, J. S. Sefadi, Morphology and Properties of Electrospun PCL and Its Composites for Medical Applications: A Mini Review. Applied Sciences 9, 2205 (2019).

10. A. K. Sharma, R. Bhandari, A. Aherwar, R. Rimašauskienė, Matrix materials used in composites: A comprehensive study. Materials Today: Proceedings 21, 1559– 1562 (2020).

11. P. S. Goh, A. F. Ismail, B. C. Ng, Directional alignment of carbon nanotubes in polymer matrices: Contemporary approaches and future advances. Composites Part A: Applied Science and Manufacturing 56, 103–126 (2014).

12. Q. Wang, J. Dai, W. Li, Z. Wei, J. Jiang, The effects of CNT alignment on electrical conductivity and mechanical properties of SWNT/epoxy nanocomposites. Composites Science and Technology 68, 1644–1648 (2008).

13. H. Zhang, L. Zhu, F. Zhang, M. Yang, Effect of Fiber Content and Alignment on the Mechanical Properties of 3D Printing Cementitious Composites. Materials 14, 2223 (2021).

14. Y. Yang, et al., Electrically assisted 3D printing of nacre-inspired structures with self-sensing capability. Science Advances 5, eaau9490 (2019).

15. D. Kokkinis, M. Schaffner, A. R. Studart, Multimaterial magnetically assisted 3D printing of composite materials. Nat Commun 6, 8643 (2015).

16. Y. Ma, et al., Bioinspired composites reinforced with ordered steel fibers produced via a magnetically assisted 3D printing process. J Mater Sci 55, 15510–15522 (2020).

17. D. S. Melchert, et al., Flexible Conductive Composites with Programmed Electrical Anisotropy Using Acoustophoresis. Advanced Materials Technologies 4, 1900586 (2019).

18. D. E. Yunus, S. Sohrabi, R. He, W. Shi, Y. Liu, Acoustic patterning for 3D embedded electrically conductive wire in stereolithography. J. Micromech. Microeng. 27, 045016 (2017).

19. T. Ma, et al., Rheological behavior and particle alignment of cellulose nanocrystal and its composite hydrogels during 3D printing. Carbohydrate Polymers 253, 117217 (2021).

20. T. Börzsönyi, et al., Orientational Order and Alignment of Elongated Particles Induced by Shear. Phys. Rev. Lett. 108, 228302 (2012).

21. S. Behr, U. Vainio, M. Müller, A. Schreyer, G. A. Schneider, Large-scale parallel alignment of platelet-shaped particles through gravitational sedimentation. Sci Rep 5, 9984 (2015).

22. R. Sharma, C. Y. Lee, J. H. Choi, K. Chen, M. S. Strano, Nanometer Positioning, Parallel Alignment, and Placement of Single Anisotropic Nanoparticles Using Hydrodynamic Forces in Cylindrical Droplets. Nano Lett. 7, 2693–2700 (2007).

23. Y. Wu, R. Chattaraj, Y. Ren, H. Jiang, D. Lee, Label-Free Multitarget Separation of Particles and Cells under Flow Using Acoustic, Electrophoretic, and Hydrodynamic Forces. Anal. Chem. 93, 7635–7646 (2021).

24. Z. Zhang, A. Sukhov, J. Harting, P. Malgaretti, D. Ahmed, Rolling microswarms along acoustic virtual walls. Nat Commun 13, 7347 (2022).

25. T. M. Llewellyn-Jones, B. W. Drinkwater, R. S. Trask, 3D printed components with ultrasonically arranged microscale structure. Smart Mater. Struct. 25, 02LT01 (2016).

26. D. Kam, et al., Wood Warping Composite by 3D Printing. Polymers 14, 733 (2022).

27. A. J. Pascall, et al., Light-Directed Electrophoretic Deposition: A New Additive Manufacturing Technique for Arbitrarily Patterned 3D Composites. Advanced Materials 26, 2252–2256 (2014).

28. I. Liashenko, J. Rosell-Llompart, A. Cabot, Ultrafast 3D printing with submicrometer features using electrostatic jet deflection. Nat Commun 11, 753 (2020).

29. M. Kunitski, et al., Double-slit photoelectron interference in strong-field ionization of the neon dimer. Nat Commun 10, 1 (2019).

30. P. Romano, H. Fabritius, D. Raabe, The exoskeleton of the lobster Homarus americanus as an example of a smart anisotropic biological material. Acta Biomaterialia 3, 301–309 (2007).

31. J. Li, X. Liang, F. Liou, J. Park, Macro-/Micro-Controlled 3D Lithium-Ion Batteries via Additive Manufacturing and Electric Field Processing. Sci Rep 8, 1846 (2018).

32. Y. Yang, et al., Biomimetic Anisotropic Reinforcement Architectures by Electrically Assisted Nanocomposite 3D Printing. Advanced Materials 29, 1605750 (2017).

33. D. Han, C. Yang, N. X. Fang, H. Lee, Rapid multi-material 3D printing with projection micro-stereolithography using dynamic fluidic control. Additive Manufacturing 27, 606–615 (2019).

34. J. J. Martin, B. E. Fiore, R. M. Erb, Designing bioinspired composite reinforcement architectures via 3D magnetic printing. Nat Commun 6, 8641 (2015).

35. R. M. Erb, D. S. Sebba, A. A. Lazarides, B. B. Yellen, Magnetic field induced concentration gradients in magnetic nanoparticle suspensions: Theory and experiment. Journal of Applied Physics 103, 063916 (2008).

36. X. Wang, M. Jiang, Z. Zhou, J. Gou, D. Hui, 3D printing of polymer matrix composites: A review and prospective. Composites Part B: Engineering 110, 442– 458 (2017).

37. H. Song, et al., Inkjet printing of magnetic materials with aligned anisotropy. Journal of Applied Physics 115, 17E308 (2014).

38. T. Nakamoto, S. Kojima, Layered Thin Film Micro Parts Reinforced with Aligned Short Fibers in Laser Stereolithography by Applying Magnetic Field. *Journal of Advanced Mechanical Design*, Systems, and Manufacturing 6, 849–858 (2012).

39. Y. Wang, et al., Acoustic-assisted 3D printing based on acoustofluidic microparticles patterning for conductive polymer composites fabrication. Additive Manufacturing 60, 103247 (2022).

40. J. Durrer, et al., A robot-assisted acoustofluidic end effector. Nat Commun 13, 6370 (2022).

41. K. Johnson, D. Melchert, D. S. Gianola, M. Begley, T. R. Ray, Recent progress in acoustic field-assisted 3D-printing of functional composite materials. MRS Advances 6, 636–643 (2021).

42. L. Lu, X. Tang, S. Hu, Y. Pan, Acoustic Field-Assisted Particle Patterning for Smart Polymer Composite Fabrication in Stereolithography. 3D Printing and Additive Manufacturing 5, 151–159 (2018).

43. M.-S. Scholz, B. W. Drinkwater, R. S. Trask, Ultrasonic assembly of anisotropic short fibre reinforced composites. Ultrasonics 54, 1015–1019 (2014).

44. X. Li, K. M. Lim, W. Zhai, A novel class of bioinspired composite via ultrasound-assisted directed self-assembly digital light 3D printing. Applied Materials Today 26, 101388 (2022).

45. B. E. Kelly, et al., Volumetric additive manufacturing via tomographic reconstruction. Science 363, 1075–1079 (2019).

46. P. N. Bernal, et al., Volumetric Bioprinting of Complex Living-Tissue Constructs within Seconds. Advanced Materials 31, 1904209 (2019).

47. M. Aliabouzar, et al., Micropatterning of acoustic droplet vaporization in acoustically-responsive scaffolds using extrusion-based bioprinting. Bioprinting 25, e00188 (2022).

48. Y. Sriphutkiat, S. Kasetsirikul, D. Ketpun, Y. Zhou, Cell alignment and accumulation using acoustic nozzle for bioprinting. Sci Rep 9, 17774 (2019).

49. R. R. Collino, et al., Deposition of ordered two-phase materials using microfluidic print nozzles with acoustic focusing. Extreme Mechanics Letters 8, 96–106 (2016).

50. R. R. Collino, et al., Scaling relationships for acoustic control of two-phase microstructures during direct-write printing. Materials Research Letters 6, 191– 198 (2018).

51. K. Niendorf, B. Raeymaekers, Using supervised machine learning methods to predict microfiber alignment and electrical conductivity of polymer matrix composite materials fabricated with ultrasound directed self-assembly and stereolithography. Computational Materials Science 206, 111233 (2022).

52. A. A. Doinikov, Acoustic radiation pressure on a compressible sphere in a viscous fluid. Journal of Fluid Mechanics 267, 1–22 (1994).

53. Z. Meng, X. Zhai, J. Wei, Z. Wang, H. Wu, Absolute Measurement of the Refractive Index of Water by a Mode-Locked Laser at 518 nm. Sensors 18, 1143 (2018).

54. H. Bruus, Acoustofluidics 7: The acoustic radiation force on small particles. Lab on a Chip 12, 1014–1021 (2012).

55. M. Xie, et al., Volumetric additive manufacturing of pristine silk-based (bio)inks. Nat Commun 14, 210 (2023).

56. C. E. Garciamendez-Mijares, P. Agrawal, G. García Martínez, E. Cervantes Juarez, Y. S. Zhang, State-of-art affordable bioprinters: A guide for the DiY community. Applied Physics Reviews 8, 031312 (2021).

57. D. Loterie, P. Delrot, C. Moser, High-resolution tomographic volumetric additive manufacturing. Nat Commun 11, 852 (2020).

58. Thingiverse.com, Thingiverse - Digital Designs for Physical Objects (August 4, 2023).

## SI References

1. When it Comes to Glass, Don’t “Glaze-over” Acoustics! Acoustics First BLOG (2022) (February 21, 2023).

2. J.-M. Wu, H.-L. Huang, Microwave properties of zinc, barium and lead borosilicate glasses. Journal of Non-Crystalline Solids 260, 116–124 (1999).

3. J. van Deventer, J. Delsing, Thermostatic and dynamic performance of an ultrasonic density probe. IEEE Transactions on Ultrasonics, Ferroelectrics, and Frequency Control 48, 675–682 (2001).

4. M. Schwartz, Reflection, Transmission and Impedance.

5. Waves at interfaces — GPG 0.0.1 documentation (February 21, 2023).

6. J. Shi, et al., Acoustic tweezers: patterning cells and microparticles using standing surface acoustic waves (SSAW). Lab Chip 9, 2890–2895 (2009).

7. A. A. Doinikov, Acoustic radiation pressure on a compressible sphere in a viscous fluid. Journal of Fluid Mechanics 267, 1–22 (1994).

8. Z. Zhang, A. Sukhov, J. Harting, P. Malgaretti, D. Ahmed, Rolling microswarms along acoustic virtual walls. Nat Commun 13, 7347 (2022).

9. Stainless Steel (Non-Magnetic) Microspheres 7.8g/cc - 1um to 1200um (1.2mm). Cospheric (February 21, 2023).

10. P. W. Bridgman, The Compressibility of Thirty Metals as a Function of Pressure and Temperature. Proceedings of the American Academy of Arts and Sciences 58, 165–242 (1923).

